# Helical Repeat Protein mRSiC is Required for rRNA Fragment Accumulation in *T. gondii* Mitochondria

**DOI:** 10.64898/2026.01.22.701003

**Authors:** Nikiforos Drakoulis, Sascha Maschmann, Zala Gluhic, Alea Boujianna Blum, Arne Hillebrand, Alexandra Possling, Victor Flores, Ross Waller, Giel van Dooren, Christian Schmitz-Linneweber, Sabrina Tetzlaff

## Abstract

Myzozoans, including apicomplexan parasites, possess highly reduced and unusual mitochondrial genomes that encode two to three respiratory chain subunits and extensively fragmented rRNAs forming a divergent mitoribosome. Heptatricopeptide repeat (HPR) proteins are expanded in myzozoans, with some functioning as mitoribosomal constitutes. Additional HPR proteins have been proposed to function in post-transcriptional processes of mitochondrial gene expression, though their specific roles remain unexplored. We present a phylogenetic analysis of HPR proteins that reveals extensive lineage-specific expansions, consistent with diversifications of mitochondrial gene expression systems within myzozoans. We further characterize mRSiC, a coccidian-specific HPR found in *Toxoplasma gondii* and its closest relatives. Using sRNA sequencing and RNA gel blot analyses, we show that mRSiC is required for the stabilization of two coccidian-specific mitochondrial sRNAs, RNA33 and RNA42. Loss of mRSiC causes rapid depletion of these RNAs, followed by secondary reductions of rRNA fragments, defects in the respiratory chain, and impaired parasite proliferation. Together, these findings highlight the importance of HPRs in apicomplexan mitochondrial gene expression and illustrate how lineage-specific RNA-binding factors support highly derived mitochondrial expression systems.

## Introduction

Mitochondrial genomes in Myzozoa display a remarkable degree of reduction and structural idiosyncrasy. A defining feature of this system is that the ribosomal RNAs (rRNAs) are not encoded and transcribed as continuous molecules but instead as a multitude of small fragments (Waller and Jackson, 2009; Feagin *et al*., 2012; Oborník and Lukeš, 2015; Tetzlaff *et al*., 2024; Wang *et al*., 2024; Shikha *et al*., 2025). The *Toxoplasma gondii* mitochondrial genome (mitogenome) represents one of the most extreme cases characterized to date. It is highly fragmented and organized into extensive - but non-random - repeated sequence blocks that encode only a minimal set of proteins and ribosomal RNAs (rRNAs) (Berná *et al*., 2021a; Namasivayam *et al*., 2021; Tetzlaff *et al*., 2024).

This extreme mitogenome fragmentation and the associated recombination of sequence blocks are not universal within apicomplexans but appear to be evolutionarily restricted. The phenomenon is characteristic of members of the Sarcocystidae, including *Toxoplasma gondii*, *Neospora caninum*, and *Hammondia hammondi*, all of which possess mitochondrial genomes composed of short sequence blocks that recombine to generate diverse and dynamic genomic configurations (Berná *et al*., 2021a; Namasivayam *et al*., 2021; Tetzlaff *et al*., 2024). In contrast, other apicomplexans such as *Plasmodium falciparum*, *Babesia microti*, and *Theileria parva* maintain compact, nonrecombining linear mitochondrial genomes, typically organized as monomers or tandemly repeated units of ∼6 kb to ∼12 kb that encode the same minimal gene complement (Kairo *et al*., 1994; Feagin *et al*., 2012; Hikosaka *et al*., 2012). These contrasts suggest that block-based recombination and extensive genomic plasticity evolved secondarily within the Sarcocystidae, representing a derived and lineage-specific feature.

More broadly, myzozoan mitochondrial gene expression systems exhibit fascinating lineage-specific innovations, including ribosomal frameshifting in *Perkinsus* (Gornik *et al*., 2022), and extensive RNA editing and trans-splicing in dinoflagellates (Waller and Jackson, 2009). Together these examples illustrate how unconventional mitochondrial genome architectures in myzozoans are accompanied by equally unusual gene expression mechanisms.

Following the initial demonstration that fragmented rRNAs are incorporated into functional mitochondrial ribosomes (mitoribosomes) (Tetzlaff *et al*., 2024), high-resolution structural analyses of the *T. gondii* mitochondrial ribosome revealed the presence of numerous distinct rRNA fragments and defined their architectural roles within the ribosome (Wang *et al*., 2024; Shikha *et al*., 2025). These studies established that fragmented rRNAs are indeed functional and are continuously required for ribosome biogenesis - and thus for mitochondrial activity. There is only very limited insight into how these rRNAs are produced. What is known is that transcription of the mitochondrial genome in apicomplexans produces longer, polycistronic precursor RNAs (Ji *et al*., 1996; Namasivayam *et al*., 2021). Central questions remain: which factors define RNA processing boundaries, and how are transcripts stabilized or degraded?

In other systems, nucleases and RNA helicases play important roles in RNA maturation, often in conjunction with RNA-binding proteins (RBPs). In particular, helical repeat proteins including PPR (pentatricopeptide repeat) have emerged as central regulators of mitochondrial RNA metabolism. Acting as “roadblocks” to exonucleases, such proteins can delimit transcript ends and contribute to RNA processing and stabilization, as shown extensively in plants and algae (Pfalz *et al*., 2009; Zhelyazkova *et al*., 2012; Wang *et al*., 2015; Small *et al*., 2023).

In Apicomplexa, the heptatricopeptide repeat (HPR) protein family, a close relative of algae octatricopeptide proteins, was identified as a set of ∼30 HPR proteins in different apicomplexan genomes, including *Toxoplasma gondii* (Hillebrand *et al*., 2018). These proteins are characterized by tandem arrays of helical repeats with 37 amino acid periodicity, forming superhelical scaffolds that are predicted to bind RNA in a sequence-specific manner, just like other helical repeat proteins. Phylogenetic analyses revealed that - while present across eukaryotes (Boehm *et al*., 2016; Ohkubo *et al*., 2021) - HPR proteins are expanded in myzozoans, suggesting an adaptation of RNA metabolism unique to this lineage (Hillebrand *et al*., 2018). Several HPR proteins localize to the mitochondrion (Hillebrand *et al*., 2018; Hollin *et al*., 2022; Wang *et al*., 2024), consistent with a role in mitochondrial RNA biology. Based on analogy to PPR proteins in plants and algae, HPR proteins were proposed to contribute to RNA stabilization, processing, and translation, yet experimental evidence for their molecular functions has remained limited. Five *T. gondii* HPR proteins have been described to associate with the mitochondrial ribosome and form permanent subunits of the mitoribosome (Wang *et al*., 2024; Shikha *et al*., 2025). Furthermore, HPR proteins have been suggested to function in RNA processing, stabilization, or modification of mitochondrial rRNAs and tRNAs (Wang *et al*., 2024).

Here, we present a molecular analysis of mRSiC, a member of the HPR family in *Toxoplasma gondii*. By combining biochemical, genetic, and computational approaches, we dissected the role of mRSiC in mitochondrial RNA metabolism, gaining the first insights into how apicomplexan parasites organize and control their highly unconventional mitochondrial transcriptomes.

## Results

### A survey of HPR proteins across 57 genomes reveals lineage-specific expansions within Alveolata

Helical repeat proteins represent a large and dynamically evolving protein superfamily, characterized by tandem arrays of α-helical motifs. These proteins frequently arise through recombination and duplication events, leading to lineage-specific expansions and the emergence of new functions. For example, the evolution of RNA editing in land plants is thought to have been enabled by the emergence of a novel subclass of pentatricopeptide repeat (PPR) proteins (O’Toole *et al*., 2008; Fujii and Small, 2011). We hypothesized that the unusual mitochondrial genome organization in *T. gondii* and related lineages may have been accompanied by the evolution of new helical repeat proteins, specifically HPR proteins, that facilitate RNA stabilization or processing in this highly derived mitochondrial system. To explore this possibility, we searched for evolutionarily “young” HPR proteins that may have arisen within the Apicomplexa.

Previous surveys of apicomplexan helical repeat proteins identified HPRs using motifs derived from a limited set of species (*Plasmodium falciparum, Plasmodium berghei*, and *Toxoplasma gondii*), resulting in an incomplete picture of their evolutionary diversity (Hillebrand *et al*., 2018). To achieve broader coverage, we expanded this analysis by constructing a new HPR consensus motif based on a larger and more representative set of alveolates, including members of the Chromerida, Perkinsida, Dinoflagellata, and Ciliata. We assembled a training alignment from 187 HPR candidates identified in our previous study (Hillebrand *et al*., 2018) and used it to build a profile Hidden Markov Model (HMM). This resulted in a 37–amino-acid motif predicted to form a helix–turn–helix (HTH) structure (Fig 1A). Notably, the HMM-derived motif is highly similar to the PWM-based motif from our earlier study (Hillebrand *et al*., 2018), indicating that both independent approaches converge on a robust consensus for HPR detection.

**Figure 1.**
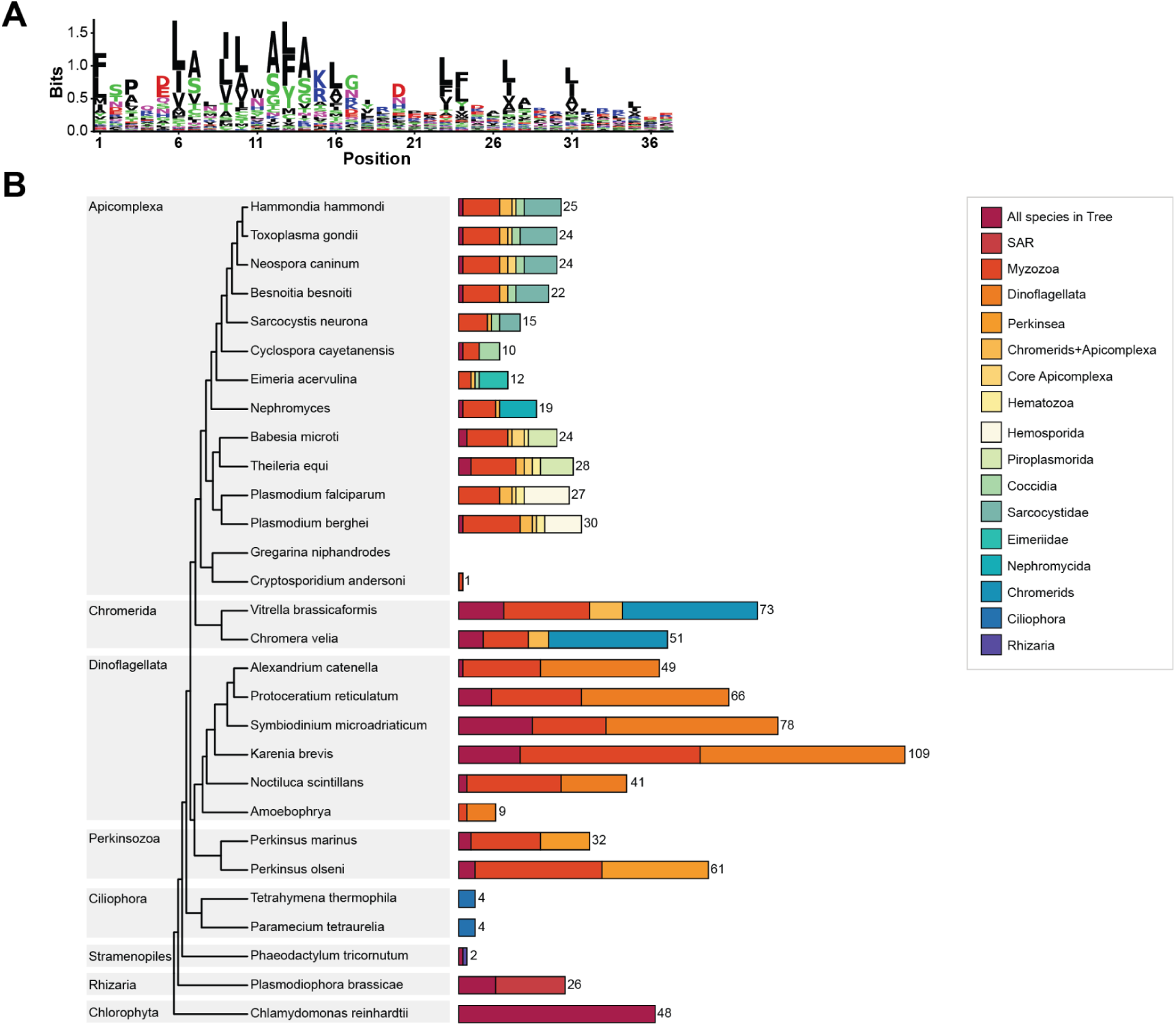
A multi-lineage survey uncovers ancient and lineage-specific expansions of HPR proteins. **(A)** A 37-aa helix–turn–helix HPR motif was derived by building an HMM from a curated multi-alveolate alignment of 137 HPR proteins reported by (Hillebrand *et al*., 2018). The bit logo illustrates residue preferences used for all motif searches in this study. **(B)** Species tree with MRCA-resolved HPR repertoires. A schematic alveolate species tree curated from published literature was used as the reference framework. HPR proteins detected by the alveolate HMM were grouped using OrthoFinder, and each orthogroup was assigned to the most recent common ancestor (MRCA) of its member taxa. For each species, a horizontal stacked bar shows the number of HPR proteins originating at each MRCA node. Colors correspond to evolutionary node groups as specified in the legend. SAR = Stramenopiles + Alveolates + Rhizaria

Using the alveolate HPR HMM, we scanned proteomes from 57 species spanning a *Chlamydomonas* outgroup, dinoflagellates, perkinsids, chromerids, apicomplexans, and additional alveolate taxa. This survey identified 2,433 HPR proteins containing at least two tandem repeats, with a mean of approximately six repeats per protein (Fig S1, Table S2). For comparison, plant PPR proteins typically comprise long arrays of 10 to 20 (or more) ∼ 35-aa repeats that together form a superhelical RNA-binding surface with residue-level specificity (Small and Peeters, 2000; Barkan *et al*., 2012). The substantially shorter repeat tracts we observe in alveolate HPRs may reflect either recognition of shorter RNA elements or a divergent mode of RNA binding involving a stronger contribution from non-HPR regions.

We then proceeded to group the identified HPR proteins into orthogroups and mapped them onto a schematic species phylogeny, revealing both broad conservation and pronounced lineage-specific variation (Fig 1B, Table S2). For the species included in our earlier search using the more species-restricted motif (Hillebrand *et al*., 2018) the number of HPR proteins identified closely matched those recovered with our new alveolate HPR consensus (Fig 1B), underscoring the robustness of the approach.

Our analysis corroborates the expanded repertoire of HPR proteins in Myzozoa and in other species that possess fragmented rather than contiguous mitochondrial rRNAs, such as *Chlamydomonas* (Fig 1B). Of note, our alveolate HPR motif is able to detect proteins previously characterized as OPRs in *Chlamydomonas* (Fig 1B) -though not the full complement reported by Cattelin *et al*., (preprint, 2025). To contextualize this, we compared our alveolate HPR consensus motif with a motif trained on chlorophyte OPR sequences (preprint: Cattelin *et al*., 2025). When contrasting the residue preferences of the two motifs, we observe clear similarities in both length and overall compositional bias (Data not shown). Nevertheless, the alveolate HPR motif performs better at capturing helical repeat–containing proteins within alveolate lineages than the chlorophyte OPR motif (Data not shown). Taken together, these observations imply that the alveolate HPR architecture is not an entirely novel invention, but rather a clade-specific adaptation within a broader family of conserved helical repeat scaffolds. In line with this, HPR proteins like OPR proteins often show shorter helical repeat tracts and can carry additional FAST kinase–like or RAP domains associated with RNA metabolism (Simarro *et al*., 2010; Eberhard *et al*., 2011; Hammani *et al*., 2014). Given these sequence and structural similarities, further features described for OPRs may also apply to alveolate HPR proteins. This includes the high sequence degeneracy described for OPR motifs (Rahire *et al*., 2012; Hammani *et al*., 2014).

Remarkably, our phylogenetic analysis revealed lineage-specific expansions of the HPR family-for example, in Chromerida and in Dinoflagellata (blue and orange bars, respectively, in Fig 1B). Of particular relevance, 9 HPR proteins were found to be specific to the Sarcocystidae with no ortholog detected in the closely related Eimeridae (blue-green bars in Fig 1B). The concurrent appearance of these HPR expansions with the emergence of the highly repetitive, sequence-block–based mitochondrial genome organization in the Sarcocystidae suggests that these proteins may have evolved to support gene expression in the context of this dynamic genome architecture.

### mRSiC as a representative of an evolutionarily “young” HPR orthogroup

We next examined the *T. gondii* HPR repertoire in more detail by mapping the inferred origin of each orthogroup onto the species tree. For every *T. gondii* HPR-containing orthogroup, we recorded the most recent common ancestor (MRCA) of all member taxa and used this MRCA to assign an “age” to the corresponding *T. gondii* protein. The resulting ladder plot (Fig 2A) displays individual *T. gondii* HPR proteins on the x-axis and the lineage at which their orthogroup first emerged on the y-axis. This analysis revealed a layered history: an older core of HPR proteins with orthologs extending to *Chlamydomonas* or deep-branching alveolates and myzozoans, and a conspicuous set of 11 “young” proteins whose orthogroups are restricted to Coccidia and, within that group, often to Toxoplasmatinae (Fig 2A). From this set, we selected one representative gene, TGME49_237530, which belongs to an orthogroup traceable from *Toxoplasma* through *Hammondia*, *Neospora*, *Besnoitia* and *Sarcocystis* up to *Cyclospora*, but not beyond Coccidia (Fig 2A). We hereafter refer to the protein encoded by TGME49_237530 as mitochondrial RNA Stabilizer in Coccidia (mRSiC), based on its proposed function described later in this study. Genome-wide CRISPR knockout screens indicated that mRSiC contributes moderately to parasite fitness (Sidik *et al*., 2016). Because strong knockouts often lead to pleiotropic growth defects and secondary effects, we reasoned that analyzing a gene with a milder phenotype would be more suitable for dissecting the molecular functions of HPR proteins.

**Figure 2.**
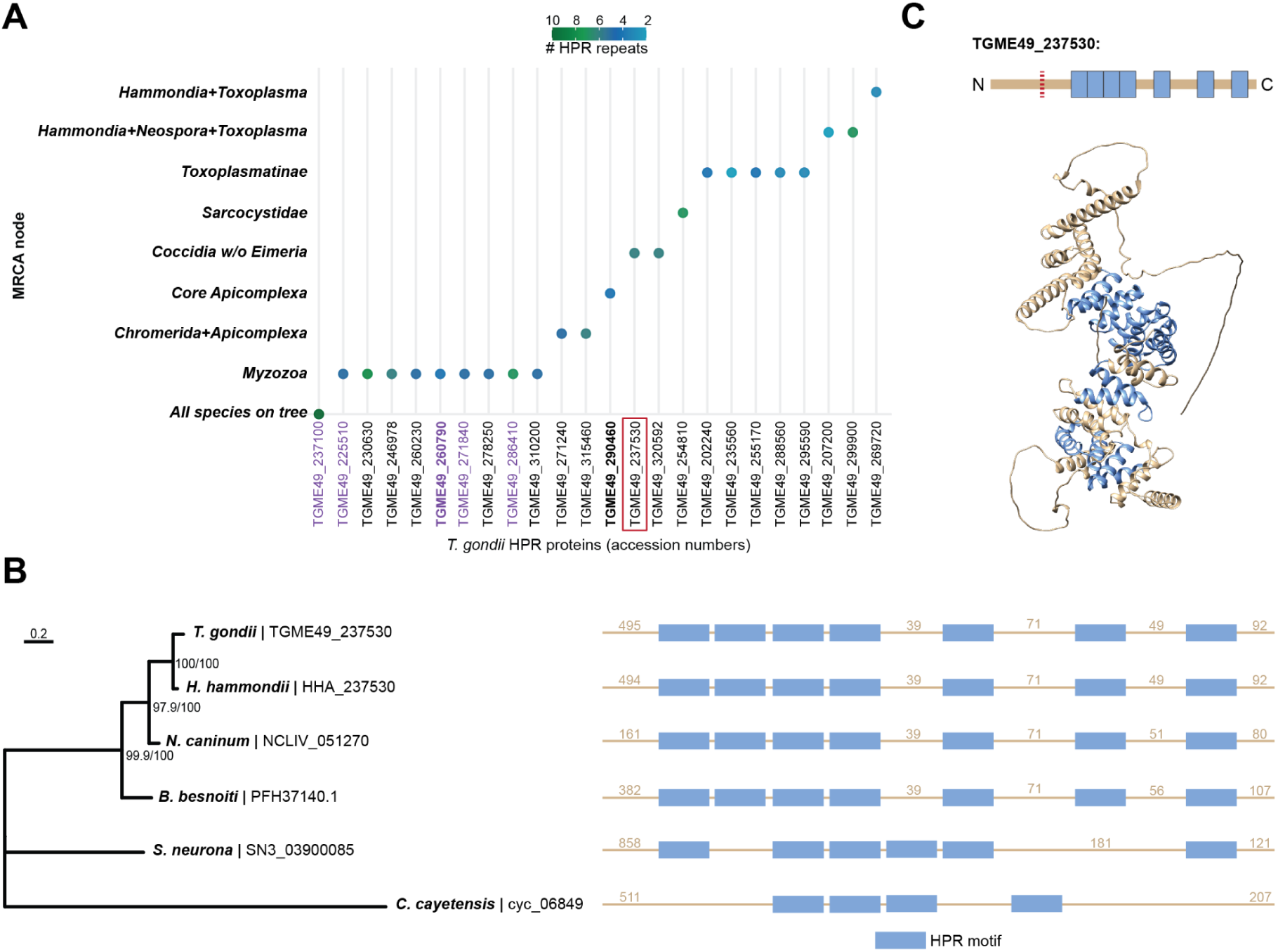
mRSiC defines an evolutionarily young HPR lineage. **(A)** *Toxoplasma gondii* HPR MRCA profile. For each *T. gondii* HPR protein, the MRCA of its orthogroup was mapped onto the species tree, generating a ladder plot that displays the evolutionary depth at which each protein first appeared. Colors denote the number of HPR repeats detected in each protein. The protein selected for functional characterization in this study, TGME49_237530, is highlighted by a red rectangle. Proteins in bold also contain a RAP domain. Proteins in purple are part of the mitochondrial ribosome. **(B)** Left panel: Maximum-likelihood phylogeny of mRSiC orthologs across Coccidia, inferred with IQ-TREE using the best-fit substitution model; branch support values are shown at internal nodes. Right panel: Motif diagram of mRSiC orthologs shown in the left panel, indicating the positions of HPR repeats (light blue) detected by the alveolate HMM. All Coccidian orthologs (*Toxoplasma, Hammondia, Neospora, Besnoitia, Sarcocystis*) share a conserved core tract of four tandem repeats, with additional repeats positioned outside the main array. Non-HPR motif regions are shown in beige and are not drawn to scale. **(C)** Top panel: Schematic representation of *T. gondii* mRSiC (TGME49_237530) with HPR motifs shown in light blue and the remaining protein regions in beige. The red dotted line marks the N-terminal segment that was omitted in the AlphaFold visualization shown below. Bottom panel: AlphaFold model of *T. gondii* mRSiC. The N-terminal 273 aa region indicated in the upper panel (red dotted line) is predicted to be unstructured and was removed from the model to simplify visualization of the HPR tract. HPR repeats identified by the alveolate HMM are highlighted in light blue. The central protein tract folds into a continuous solenoid of helix–turn–helix hairpins, several adjacent helical hairpins with similar geometry are visible despite not being detected as HPR motifs by the HMM.

Motif diagrams for *T. gondii* mRSiC and its orthologs revealed a conserved core tract of four tandem HPR repeats and additional repeats that are partially separated from the main array (Fig 2B). The positions of these repeats are closely aligned across *Toxoplasma*, *Hammondia*, *Neospora* and *Besnoitia*, consistent with the strong branch support observed in the corresponding protein phylogeny (Fig 2B).

AlphaFold models further support this picture (Fig 2C). In *T. gondii* mRSiC, the central HPR tract is predicted to form an extended solenoid of stacked helix–turn–helix hairpins, while the N- and C-terminal regions are more flexible (Fig 2C). The models also indicate additional short helical hairpins adjacent to, but not identified as, HPR repeats by our HMM-based pipeline. These segments share the same overall HTH geometry but probably lack several of the strongly conserved positions that define the alveolate HPR consensus, thus remaining undetected by our search. In practical terms, this implies that the RNA-binding region of mRSiC may extend beyond the repeats that are strictly recognized by our profile.

Taken together, the evolutionary origin, repeat architecture, and mild fitness phenotype highlight mRSiC as an attractive candidate for investigating the function of Coccidian-specific HPR proteins in *T. gondii* mitochondrial RNA metabolism.

### mRSiC is a mitochondrial protein important for parasite growth

mRSiC was computationally predicted to localize to the *T. gondii* mitochondrion (Claros and Vincens, 1996). To test this experimentally, we introduced a C-terminal triple FLAG epitope at the endogenous locus, generating mRSiC-FLAG parasites (Fig S2A). Immunoblotting confirmed expression of the tagged protein, but the FLAG signal was detected at ∼80 kDa rather than the predicted 110 kDa for full-length mRSiC (Fig 3A). This shift in apparent molecular weight may reflect the removal of a long N-terminal sequence as part of the import process into mitochondria, similar to what was reported for *T. gondii* AP2 domain-containing mitoribosomal proteins (Wang *et al*., 2024). The ∼30 kDa N-terminal region of mRSiC is unstructured (Fig 2C) and might get removed during or after mitochondrial signal peptide cleavage, leaving the ∼80 kDa helical repeat tract. To determine the subcellular localization of mRSiC-FLAG, immunofluorescence assays were carried out using anti-FLAG antibodies together with staining for Tom40, a marker of the mitochondrial outer membrane. The FLAG signal co-localized with Tom40 (Fig 3B), confirming that mRSiC is targeted to mitochondria.

**Figure 3.**
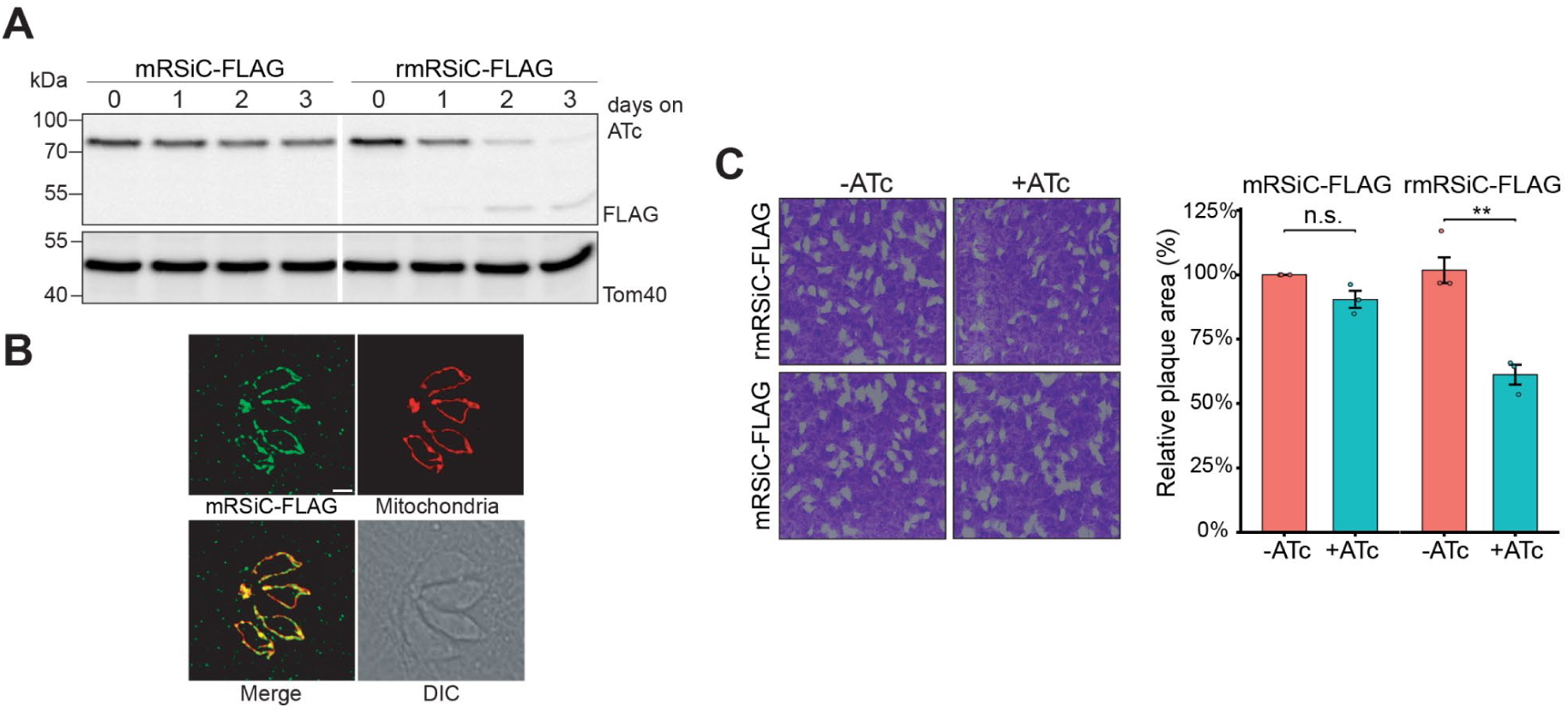
mRSiC localizes to *T. gondii* mitochondria and contributes to efficient tachyzoite proliferation. **(A)** Immunoblot analysis of rmRSiC-FLAG parasites as well as mRSiC-FLAG parasites cultured in the absence (0) or presence of anhydrotetracycline (ATc) for 1–3 days. Samples were separated by SDS–PAGE and probed with anti-FLAG and anti-TgTom40 (loading control) antibodies, showing progressive depletion of mRSiC upon ATc treatment in the rmRSiC-FLAG strain. White lines indicate where lanes not relevant for this analysis were removed. **(B)** Immunofluorescence assay of a four-cell vacuole of mRSiC-FLAG parasites stained with anti-FLAG (green) and anti-TgTom40 (red) antibodies, showing co-localization of mRSiC-FLAG with the mitochondrial outer membrane marker Tom40. Scale bar, 2 µm. **(C)** Left panel: Plaque assays assessing proliferation of mRSiC-FLAG and rmRSiC-FLAG parasites cultured with or without ATc for 7 days. Images represent one of three independent experiments. Right panel: Quantification of plaque area from three biological replicates. Data show mean ± SEM; significance determined by Welch’s t-test (–ATc vs. +ATc) for each genotype (n.s., not significant; **p < 0.01).

To investigate the function of mRSiC, we generated a conditional knockdown line in the mRSiC-FLAG background by replacing the endogenous mRSiC promoter with an anhydrotetracycline (ATc)-regulatable promoter (Fig S2B). We termed the generated line rmRSiC-FLAG and monitored mRSiC-FLAG protein levels following ATc treatment by SDS–PAGE and immunoblotting. Protein abundance was reduced after one day of ATc exposure and reached the detection limit after three days (Fig 3A, S3). In contrast, protein levels in the parental mRSiC-FLAG line remained unchanged under the same conditions (Fig 3A, S3). Notably, a smaller ∼50 kDa band appeared during knockdown, potentially corresponding to a C-terminal degradation product of mRSiC-FLAG (Fig 3A). These results demonstrate that mRSiC expression can be effectively downregulated using the ATc-regulatable TET-Off system.

Previous genome-wide CRISPR–Cas9 fitness screens identified mRSiC as a mildly counter-selected gene (phenotype score −2.53; (Sidik *et al*., 2016), suggesting a supportive role in parasite growth. To directly assess the impact of HPR on tachyzoite proliferation, mRSiC-FLAG and rmRSiC-FLAG parasites were cultured in the presence or absence of ATc, and growth was evaluated by plaque assays. Upon ATc treatment, rmRSiC-FLAG parasites displayed a significant reduction in plaque size, averaging 39.8% of the size observed in untreated controls. In contrast, the parental mRSiC-FLAG line exhibited only a minor, non-significant decrease in plaque size following ATc treatment (Fig 3C). These results indicate that mRSiC contributes to efficient tachyzoite proliferation.

### Downregulation of mRSiC diminishes Complex IV abundance

Given the dependence of electron transport chain (ETC) biogenesis on mitochondrial gene expression, and the proposed role of mRSiC in this process, we next examined the importance of mRSiC for ETC integrity. Impaired expression of a core ETC subunit -particularly those encoded by the mitochondrial genome -is known to destabilize the entire complex, leading to degradation of most, if not all, constituent subunits (Seidi *et al*., 2018; Lacombe *et al*., 2019; Hayward *et al*., 2021; Maclean *et al*., 2021; Shikha *et al*., 2022; Wang *et al*., 2024; Shikha *et al*., 2025).

To examine the impact of mRSiC depletion on ETC complex accumulation, we generated rmRSiC-FLAG parasite lines in which nuclear-encoded subunits of Complexes II, III, IV, and V were C-terminally tagged with an HA epitope: SdhB (Complex II), MPPα (Complex III), Cox2a (Complex IV), and Atpβ (Complex V) (Fig S4). Complexes II and V do not depend on mitochondrial gene expression, as all their subunits are nuclear-encoded, and thus served as controls for potential secondary effects on overall mitochondrial health.

The rmRSiC-FLAG/HA-tagged lines were cultured in the absence or presence of ATc, and protein accumulation was analyzed after three days by immunoblotting. Cox2a-HA levels were markedly reduced following ATc treatment (Fig 4A, B), whereas the levels of SdhB-HA, MPPα-HA, and Atpβ-HA remained unchanged.

**Figure 4.**
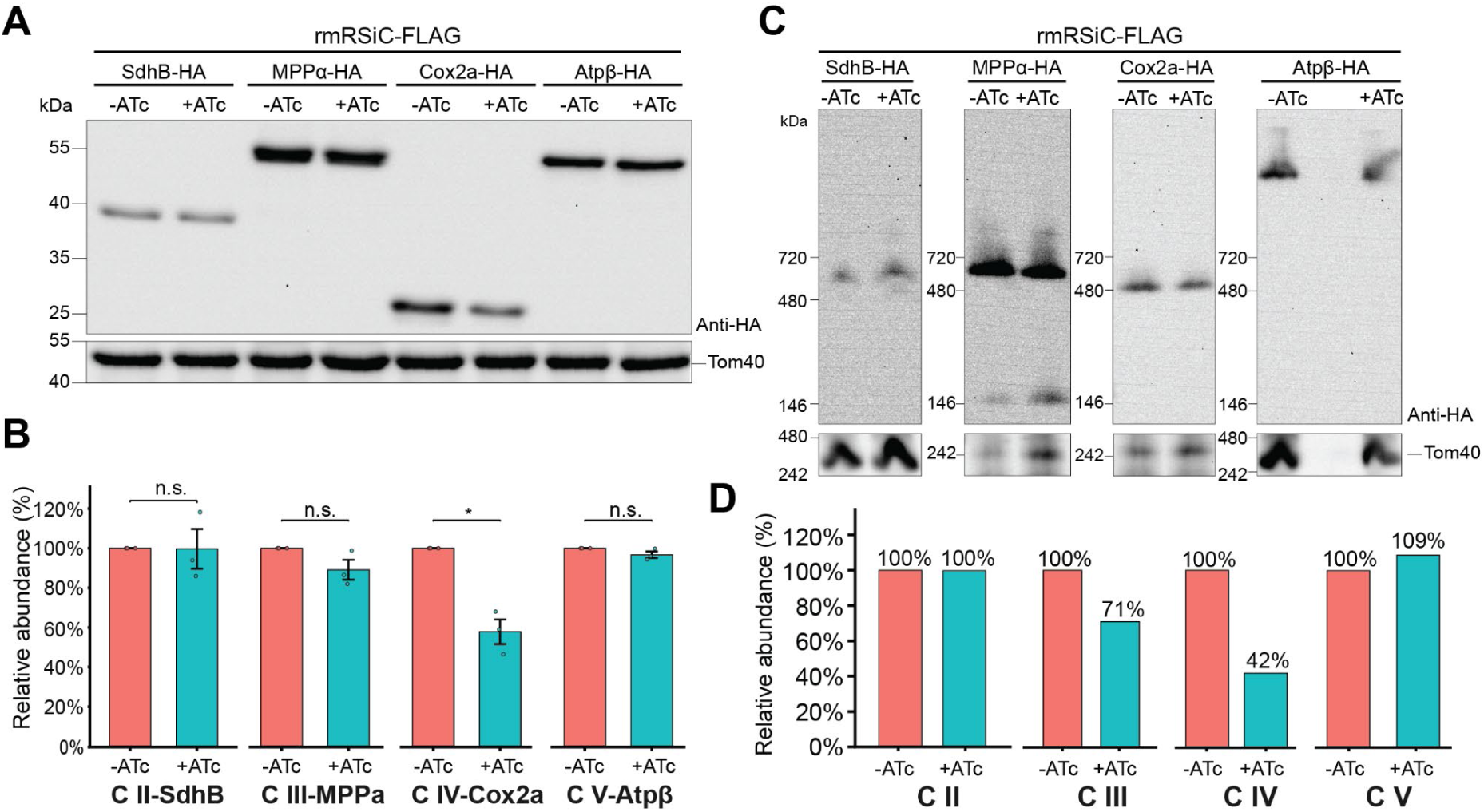
Investigation of electron transport chain (ETC) complexes following mRSiC depletion. **(A)** Immunoblot analysis of proteins extracted from rmRSiC-FLAG parasites expressing HA-tagged versions of SdhB (Complex II), MPPα (Complex III), Cox2a (Complex IV), and Atpβ (Complex V), respectively, cultured in the absence (–ATc) or presence (+ATc) of ATc for three days. Samples were separated by SDS–PAGE and probed with anti-HA and anti-TgTom40 antibodies. Calculated molecular masses: SdhB-HA - 38 kDa, MPPα-HA - 62 kDa, and Cox2a-FLAG - 28 kDa; Atpβ-FLAG - 59 kDa. Immunoblots are representative of three independent replicates. **(B)** Quantification of HA-signal intensities, normalized to the Tom40 loading control, from three independent experiments as shown in (A). Data represent mean ± SEM; statistical significance was determined by Welch’s t-test comparing –ATc and +ATc conditions for each genotype (n.s. not significant; *p < 0.01). **(C)** Blue Native-PAGE (BN-PAGE) analysis of rmRSiC-FLAG parasites expressing HA-tagged subunits of ETC Complexes II–V, cultured without (–ATc) or with (+ATc) ATc for three days. Extracts were prepared using 0.5% digitonin for Complexes II and V, and 1% (v/v) Triton X-100 for Complexes III and IV. Samples were separated by BN-PAGE and immunoblotted with anti-HA and anti-TgTom40 antibodies. **(D)** Quantification of BN-PAGE signals shown in (C), normalized to the Tom40-containing TOM complex.

To further assess complex integrity, we performed Blue Native-PAGE (BN-PAGE) followed by immunoblotting. Consistent with the reduced Cox2a-HA protein levels, normalization against the Tom40-containing translocase of the outer membrane (TOM) revealed decreased accumulation of Complex IV upon mRSiC depletion (Fig 4C, D). In addition, we observed a moderate reduction in Complex III, detected via MPPα-HA (Fig 4C, D). Previous studies have shown that MPPα protein levels are unaffected in several Complex III mutants, whereas BN-PAGE analyses of these mutants revealed that MPPα is absent from Complex III but still detectable as part of a ∼220 kDa lower-molecular-weight complex, consistent with its proposed dual role in Complex III and the *T. gondii* mitochondrial processing peptidase (Hayward *et al*., 2021). In agreement with these observations, our findings in mRSiC-depleted parasites indicate a modest reduction of MPPα-HA in Complex III while the protein remains detectable in a lower–molecular weight complex (Fig 4C, D). In contrast, downregulation of mRSiC had minimal impact on ETC Complexes II and V (Fig 4C, D).

Together, these findings demonstrate that mRSiC is specifically required for the maintenance of ETC complexes containing mitochondrially encoded subunits -Complexes III and IV, but not complexes II and V-consistent with the hypothesis that mRSiC is important for the expression of these subunits.

### Mitochondrial mRNA levels are altered during late stages of the mRSiC knockdown

Several *T. gondii* HPR proteins have recently been shown to contribute to mitochondrial function by acting as constitutive components of the mitoribosome (Wang *et al*., 2024; Shikha *et al*., 2025). Since mRSiC is not a mitoribosomal protein, its importance for ETC integrity likely reflects a role in other aspects of mitochondrial gene expression - potentially in the maturation or stabilization of mitochondrial rRNA fragments and/or mRNAs. We therefore hypothesized that mRSiC depletion might impact mitochondrial transcript levels.

To test this hypothesis, we first examined whether mRSiC knockdown affected abundance of the three mitochondrial mRNAs. We performed RNA gel blot hybridizations to analyze *coxI*, *coxIII*, and *cob* transcript levels after 1, 2, and 3 days of ATc treatment and in untreated controls. Hybridization with strand-specific RNA probes revealed a single predominant band for each mitochondrial mRNA, matching the expected open reading frame (ORF) sizes (*cob*: 1080 nt, *coxIII*: 735 nt, *coxI*: 1476 nt) reported previously (Namasivayam *et al*., 2021; Tetzlaff *et al*., 2024), Fig 5A-C). As a control we hybridized a probe against the cytosolic actin (*act1*) mRNA (Fig 5D), which is not expected to be influenced by mRSiC depletion. Depletion of mRSiC only moderately affected the levels of mitochondrial mRNAs, with changes emerging only at later stages of the knockdown (Fig 5A-C). After normalizing to *act1,* we detected an ∼1.3-fold increase in *cob* abundance after 2 days of mRSiC knockdown, which persisted after 3 days (Fig 5A, E). Similarly, *coxIII* levels increased by ∼1.25-fold after 2 days of ATc treatment, but dropped to ∼75% of the levels detected in untreated parasites after 3 days of treatment (Fig 5B, E). Changes in *coxI* levels became apparent only after 3 days of ATc treatment, displaying a ∼35% decrease compared to untreated parasites (Fig 5C, E).

**Figure 5.**
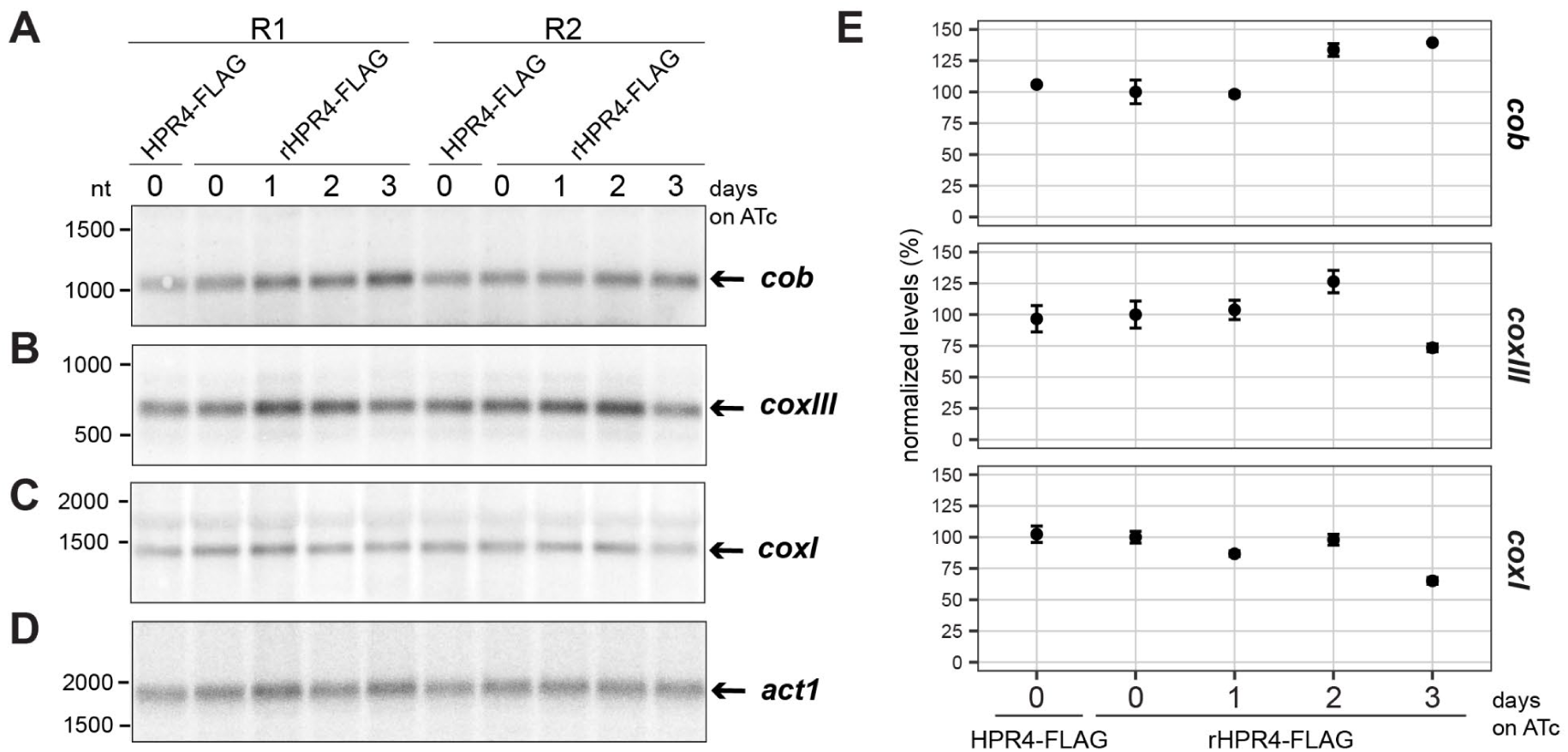
Loss of mRSiC mildly affects mitochondrial mRNA abundance. **(A)** RNA gel blot analysis of total RNA (5 µg per sample) extracted from untreated parental parasites (mRSiC-FLAG) and from rmRSiC-FLAG parasites cultured in the absence or presence of anhydrotetracycline for 1–3 days. For each condition, two independent biological replicates were analyzed (R1 and R2). The *cob* mRNA was detected with a strand-specific radiolabeled antisense probe, producing a signal at the expected size of the cob ORF (1080 nt) that showed a slight increase in intensity after 2 and 3 days of ATc treatment. **(B-D)** The blot from panel A was reprobed with probes against *coxIII* (B), *coxI* (C) and cytosolic *act1* (D) as a loading control. Between consecutive hybridizations, the blot was stripped under denaturing conditions. For each mRNA, one predominant signal at the expected size was detected (*coxIII*: 735 nt, *coxI*: 1476 nt, *act1*: 1883 nt). After 3 days of ATc treatment, a modest decrease in *coxIII* and *coxI* signal intensity was observed. **(E)** Quantification of signal intensities from panels A-C, normalized to *act1* (panel D). Data represents linearly scaled absolute mean values ± SEM from two independent repeats, with mRSiC-FLAG set to 100%.

It is conceivable that the slight decrease in *coxI* and *coxIII* transcripts could contribute to the defect in Complex IV abundance upon mRSiC depletion. However, the slight increase in *cob* transcript levels does not correspond to an increase in Complex III abundance. Given the late and relatively mild impacts on mitochondrial mRNA abundances, the observed changes may be a secondary rather than a primary effect of mRSiC depletion.

### mRSiC is required for mitochondrial rRNA fragment accumulation

Next, we specifically analyzed the levels of the multitude of *T. gondii* mitochondrial rRNA fragments in mRSiC knockdown parasites by performing a dedicated small RNA (sRNA) sequencing protocol. To enrich mitochondrial transcripts and minimize contamination with cytosolic and apicoplast RNAs, we prepared organelle-enriched fractions as described previously (Tetzlaff *et al*., 2024) and further depleted cytosolic and apicoplast rRNAs by using a customized rRNA depletion approach (Thompson *et al*., 2020). This workflow markedly increased the proportion of mitochondrial reads from less than 10% in earlier attempts (Tetzlaff *et al*., 2024) to over 25% of total reads (Fig 6A), substantially improving the representation of mitochondrial rRNA fragments in the dataset.

**Figure 6.**
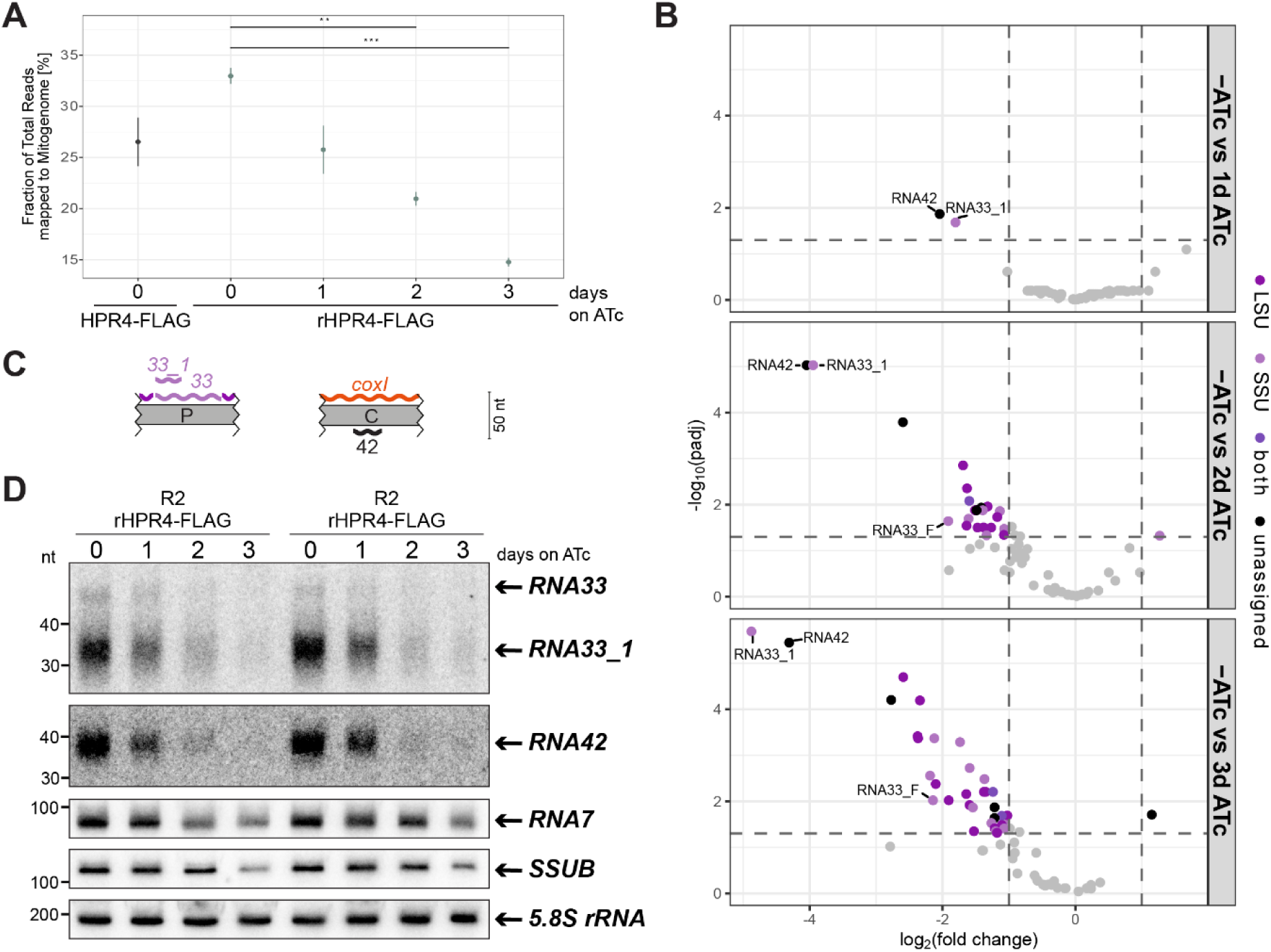
mRSiC depletion strongly reduces RNA33 and RNA42 levels during early knockdown stages and leads to a global decrease in mitochondrial rRNA accumulation. **(A)** Percentage of total reads mapping to the mitochondrial pseudogenome in mRSiC-FLAG and rmRSiC-FLAG parasites treated with ATc for 0–3 days. After 2 and 3 days of mRSiC knockdown, the proportion of mitochondrial reads is significantly reduced relative to samples without knockdown induction. Data represent mean ± SEM; n=3; statistical significance was assessed using pairwise Welch t-tests on logit-transformed values, with Benjamini–Hochberg correction for multiple testing (**p < 0.01, ***p < 0.001). **(B)** Volcano plots representing differential accumulation of mitochondrial small RNAs determined by sRNA-Sequencing, comparing untreated rmRSiC-FLAG parasites versus rmRSiC-FLAG parasites treated for 1 (upper panel), 2 (middle panel), or 3 days (lower panel) with ATc. Transcripts with log_2_(fold change) >= 1 and padj < 0.05 are color-coded according to their ribosomal subunit assignment, all remaining transcripts shown in gray. RNA42 and RNA33_1 exhibit the earliest and strongest change upon mRSiC depletion, whereas the majority of mitochondrial sRNAs are reduced only at later knockdown stages. n=3; statistical significance was assessed using edgeR quasi-likelihood F-tests with Benjamini-Hochberg-adjusted p-values. **(C)** Schematic representation of the genomic locations of RNA33_1 and RNA42 within the *T. gondii* mitogenome. Both transcripts were newly identified in this study. RNA33_1 is a truncated version of RNA33, and RNA42 encoded antisense to *coxI*. Mitochondrial sequence blocks are shown in gray, and RNAs depicted as wiggle lines, with the colors indicating their ribosomal subunit assignment (legend as in panel B). RNA33_1 was assigned based on the affiliation of full-length RNA33, although its integration into the ribosome has not been experimentally demonstrated. **(D)** Small RNA gel blot analysis of RNA (15 µg per sample) extracted from rmRSiC-FLAG parasites treated for 0-3 days with ATc using two independent biological replicates (R1 and R2) per condition. Small RNAs were detected using ^32^P-labeled oligonucleotides antisense to the transcript of interest. Between each hybridization (RNA33 → RNA42 → RNA7 → SSUB → 5.8S rRNA), probes were removed under denaturing conditions. Hybridization of the cytosolic 5.8S rRNA served as a loading control. Signals for RNA33 and RNA42 are reduced after 1 day of mRSiC depletion and reach the detection limit after 3 days. RNA7 and SSUB are affected more mildly and later during the knockdown. Signal quantification is shown in Fig S6.

We analyzed rRNA fragment accumulation in parental mRSiC-FLAG parasites and in rmRSiC-FLAG parasites that were either untreated or treated with ATc for 1, 2, or 3 days, using 3 independent replicates per condition. Given the complexity of the *T. gondii* mitochondrial genome and small RNA transcriptome, we generated a pseudogenome reference representing all two-block combinations present in the mitogenome (Namasivayam *et al*., 2021; Tetzlaff *et al*., 2024) to facilitate comprehensive mapping of mitochondrial reads. First, to assess whether mRSiC depletion causes a global change in mitochondrial gene expression, we determined the proportion of reads mapping to the mitochondrial pseudogenome relative to the total number of reads in each sRNA-seq sample. Indeed, we observed a progressive decline in the proportion of mitochondrial reads over the course of ATc treatment (Fig 6A). In untreated controls, mitochondrial reads accounted for approximately 33% of total reads, after 2 days of mRSiC knockdown this proportion significantly dropped to ∼21% and decreased further to ∼15% after three days of knockdown (Fig 6A). Given the minimal effects of mRSiC depletion on mito mRNA levels (Fig 5), this rapid and global reduction of mitochondrial reads after mRSiC depletion, indicates a central role for mRSiC in maintaining mitochondrial rRNA fragment levels. Interestingly, untreated rmRSiC-FLAG parasites showed a higher proportion of mitochondrial reads compared to the untreated parental mRSiC-FLAG line (Fig 6A). This may be related to differences in mRSiC expression between these two lines. Immunoblot analysis suggested that mRSiC expression in rmRSiC-FLAG parasites is higher under the ATc-regulatable promoter compared to endogenous expression in mRSiC-FLAG parasites (Fig 3A, S3). In line with this, the elevated mitochondrial RNA levels in the uninduced knockdown line may further support that mRSiC abundance is limiting for the accumulation or stability of a substantial portion of mitochondrial rRNA transcripts.

We next aimed to examine differential accumulation of individual mitochondrial rRNA fragments upon mRSiC knockdown induction. For this analysis, we used previous rRNA fragment annotation and nomenclature (Tetzlaff *et al*., 2024; Wang *et al*., 2024). Likely due to the improved mitochondrial coverage obtained through the rRNA depletion procedure, closer inspection of the coverage profiles of untreated samples revealed several additional fragments supported by read counts comparable to, or even exceeding, those of annotated rRNA fragments (Fig S5). In most cases, these fragments appeared to be subfragments of annotated rRNAs, often coinciding with the 5’ - or 3’ ends of rRNA fragments. They may correspond to RBP-derived footprints, which have been demonstrated to accumulate in plant organelles (Ruwe and Schmitz-Linneweber, 2012; Ruwe *et al*., 2016), and were also proposed to exist in *P. falciparum* mitochondria (Hillebrand *et al*., 2018). A further possibility is that these fragments represent hitherto undescribed rRNA fragments corresponding to unassigned leftover rRNA fragments (ulr) (Wang *et al*., 2024). Accordingly, to avoid loss of information on these additional fragments and prevent interference with the quantification of validated rRNA fragments, we expanded our annotation file by adding features for RNAs supported by defined 5’- and 3’ read-end coverage exceeding a set threshold (Fig S5).

We then proceeded to analyze differential accumulation of rRNA fragments, initially comparing untreated rmRSiC-FLAG parasites with parental mRSiC-FLAG controls. Despite the mild global increase observed in rmRSiC-FLAG parasites (Fig 6A), no significant changes in the levels of individual rRNA fragments were detected between the two lines (Table S3). Consequently, in all subsequent analyses we compared ATc-treated to untreated parasites of the rmRSiC-FLAG line. We assessed differential RNA accumulation after 1 to 3 days of mRSiC knockdown, aiming to distinguish primary effects specifically caused by mRSiC loss from secondary effects resulting from mitochondrial dysfunction. Importantly, after 1 day of ATc treatment, the abundance of only two small RNAs, RNA33_1 and RNA42, was significantly changed (Fig 6B, upper panel). The transcript levels of RNA33_1 and RNA42 dropped sharply to ∼25% and ∼20% of those in untreated parasites, respectively (Fig 6B, upper panel; Table S3), possibly reflecting diminished RNA stability as a direct consequence of mRSiC depletion. After two days of mRSiC knockdown, significant reductions were observed in 19 rRNA fragments, rising to 29 fragments after 3 days (Fig 6B, middle and lower panel; Table S3). This pattern corroborated the global - possibly secondary - rRNA decrease upon mRSiC loss inferred from mitochondrial read proportions (Fig 6A). The altered rRNA fragments were distributed across both ribosomal subunits, indicating no subunit-specific bias (Fig 6B). Importantly, reductions in the levels of the two early affected transcripts, RNA33_1 and RNA42, became progressively more pronounced after 2 and 3 days of mRSiC depletion, with the magnitude and significance of their changes clearly separating them from the cluster of other RNAs decreased at the same time points (Fig 6B; Table S3).

Neither RNA33_1 nor RNA42 are part of the published rRNA sets (Tetzlaff *et al*., 2024; Wang *et al*., 2024; Shikha *et al*., 2025). RNA33_1 is 24 nt in length and represents a 3’ truncated version of RNA33 (Fig 6C; S5), which was identified as part of the small ribosomal subunit in the structure reported by Wang *et al*. (2024), but is absent from the structure published by Shikha *et al*. (2025). Notably, full-length RNA33 ranks among the most strongly reduced transcripts after 2 and 3 days of mRSiC knockdown (Fig 6B, middle and lower panel; Table S3). By contrast, RNA42, which is 26 nt long, is not a subfragment of any previously described rRNA, but was annotated here as a discrete transcript encoded at sequence block C, antisense to the *coxI* ORF (Fig 6C; S5).

To verify the expression of RNA33_1 and RNA42 with an orthogonal method and to validate the results obtained from sRNA-Sequencing, we performed RNA gel blot hybridizations. In untreated parasites, radiolabeled oligonucleotide probes complementary to RNA33_1 and RNA42 clearly detected strong signals within size ranges that roughly aligned with the trimmed read lengths detected in sRNA-Sequencing (Fig 6D; S5). Size discrepancies (∼10 to 15 nt) likely reflect 3’ end polyadenylation, a common feature of *T. gondii* mitochondrial rRNA fragments (Shikha *et al*., 2025). The probe against RNA33_1 additionally detected a faint signal matching the size of full-length RNA33 reported by (Wang *et al*., 2024).

In line with the sRNA-sequencing results, the signals for RNA33_1 and RNA42 were already markedly reduced after 1 day of ATc treatment and underwent a dramatic further decline after 2 and 3 days, reaching levels barely above the detection limit (Fig 6D; S6). A similar trend was observed for full-length RNA33 (Fig 6D; S6). We additionally hybridized probes against the conserved rRNA fragments RNA7 and SSUB, serving as representatives of the broader set of mitochondrial rRNAs that, according to the RNA-sequencing data, were affected less strongly and only at later knockdown stages (Fig 6B, Table S3). RNA gel blot analysis confirmed these results, with signals for RNA7 and SSUB exhibiting noticeably later and milder reductions in response to mRSiC depletion compared to RNA33_1 and RNA42 (Fig 6D; S6).

### RNA33 and RNA42 are poly(A)-tailed independently of mRSiC

The size discrepancies observed for RNA33_1 and RNA42 between sRNA-Sequencing and RNA gel blotting together with the proposed importance of rRNA polyadenylation for structural integrity of the *T. gondii* mitoribosomes (Shikha *et al*., 2025), prompted us to examine the polyadenylation status of transcripts in our sRNA-sequancing dataset. Overall, we confirmed the transcript-specific polyadenylation patterns reported by Shikha *et al*. (2025) (Table S4). RNA33_1 and RNA42 were both found to be polyadenylated in untreated parental and rmRSiC-FLAG parasites (Fig S7; Table S4). For both transcripts, the length of A-tails ranged from 8 to 17 nt, with a peak at 12 nt (Fig S7; Table S4). These lengths match exactly the differences between trimmed read lengths and transcript sizes detected by RNA gel blotting (Fig 6D; S5). Moreover, the variability in A-tail length likely accounts for the diffuse, rather than discrete, signals for RNA33_1 and RNA42 in RNA gel blots (Fig 6D). Notably, in agreement with the data from Wang *et al*. (2024) and the size detected in RNA gel blot hybridization (Fig 6D), the majority of full-length RNA33 transcripts lacks A-tails (Fig S7; Table S4). Finally, we asked whether the Poly-A tail length of mitochondrial sRNAs was altered following mRSiC depletion. For the majority of mitochondrial sRNAs, the Poly-A tail length remained unchanged after mRSiC knockdown (Table S5). However, after 3 days of ATc treatment, the Poly-A tails of RNA42 and full-length RNA33 were significantly extended by ∼3–8 nt and ∼3-4 nt, respectively, relative to untreated controls (Fig S7, Table S5). These late and relatively modest effects on polyadenylation likely reflect secondary, pleiotropic consequences of mRSiC loss; nonetheless, they further support a specific contribution of mRSiC to RNA33 and RNA42 transcript fidelity.

Finally, based on the hypothesis that HPRs bind to RNA in a comparable sequence-specific manner as PPR proteins (Barkan *et al*., 2012), we searched for sequence similarities between RNA33/RNA33_1 and RNA42. Remarkably, they have perfectly overlapping sequence features at their 5’ ends (Fig S8). In particular, 10 out of the 17 first nucleotides at the 5’ end are identical. Notably, we did not find any other exact matches for the sequence elements shared between the RNA33 and RNA42 5’ ends within the *T. gondii* mitochondrial genome.

In sum, these findings demonstrate a critical role for mRSiC in sustaining the steady-state levels of a wide range of rRNA fragments in *T. gondii* mitochondria and reveal a particular importance of mRSiC for the accumulation or stability of RNA33 and RNA42.

## Discussion

In apicomplexan parasites, mitochondrial genomes and their expression are defined by a multitude of remarkable peculiarities. Mitogenomes are highly reduced and in some cases recombinant (Berná *et al*., 2021b), transcriptomes exhibit extensive sense- and antisense overlaps (Tetzlaff *et al*., 2024), and the mitoribosomes are strikingly divergent consisting of more than 40 rRNA fragments and numerous clade-specific ribosomal proteins (Wang *et al*., 2024; Shikha *et al*., 2025). Despite growing insight into these unusual and fascinating features, the mechanistic and regulatory principles governing mitochondrial gene expression in apicomplexans remain largely unresolved. Previous studies suggested that the evolution of these atypical gene expression systems was accompanied by expansions of putative RNA-binding protein families, in particular RAPs and HPRs (Hillebrand *et al*., 2018; Hollin *et al*., 2021). Here, we employed a phylogenetic approach to examine the evolution of HPR proteins across the alveolate superphylum and subsequently functionally characterized one coccidian-specific member, mRSiC, in *T. gondii*, thereby linking lineage-specific evolution to functional specialization.

### Lineage-specific HPRs in the context of intricate gene expression systems in alveolates

Examining the phylogenetic distribution of HPR proteins uncovered a small clade of evolutionarily ancient proteins conserved across nearly all surveyed alveolates, alongside a second, larger group restricted to specific lineages or species. Future studies focusing on the functional characterization of HPR proteins belonging to the ancient group may help to trace back the ancestral role of HPRs, predating the expansions and diversifications within different alveolate lineages. As mitochondrial rRNA fragmentation evolved only in certain alveolate phyla, the ancestral HPR function may be related to more generic roles in RNA metabolism, potentially akin to those of the closely related OPR proteins in Chlorophyta, which have been implicated in mRNA stabilization and translation initiation (Rahire *et al*., 2012; Wang *et al*., 2015; Chaux *et al*., 2023).

HPRs are broadly expanded in myzozoans, an alveolate clade characterized by highly reduced mitochondrial genomes and extensively fragmented rRNAs. The acquisition of HPR proteins in these organisms has been proposed to address the unique challenges associated with processing, stabilizing, and assembling these numerous rRNA fragments (Hillebrand *et al*., 2018). Interestingly, our phylogenetic analysis uncovered only a subset of HPRs that are conserved across all myzozoans, alongside a remarkable number of lineage- or species-restricted HPRs. This pattern likely reflects on the one hand, the conservation of core features of myzozoan mitochondrial gene expression systems, and on the other, substantial lineage-specific diversification and specialization (Jackson *et al*., 2007; Flegontov *et al*., 2015; Oborník and Lukeš, 2015; Shoguchi *et al*., 2015; Berná *et al*., 2021b). Conservation and divergence within the HPR family may mirror the rRNA repertoires across myzozoans. For instance, conserved HPR proteins might be required for rRNA fragments of high importance and conservation within the mitoribosome. Cryo-EM structures of the *T. gondii* mitoribosome show that, despite extensive fragmentation, the peptidyl transferase center (PTC) is universally conserved and is formed by the LSUD/E rRNA fragment interacting with several additional fragments (Wang *et al*., 2024; Shikha *et al*., 2025). Homologs of multiple PTC-associated rRNAs occur across all myzozoan phyla (Feagin *et al*., 2012; Jackson, Gornik and Waller, 2012; Flegontov *et al*., 2015; Gornik *et al*., 2022; Haro *et al*., 2024), suggesting that myzozoan-wide HPRs may act on these deeply conserved elements. In contrast, several *T. gondii* rRNA fragments lack identifiable homologs in other myzozoans, indicating sequence divergence - particularly among peripheral rRNAs - which may be associated with the emergence of clade-or species-specific HPRs. Additional diversification likely stems from marked differences in mitogenome architecture among myzozoans (Waller and Jackson, 2009; Flegontov *et al*., 2015; Berná *et al*., 2021b; Gornik *et al*., 2022), implying corresponding differences in expression mechanisms and in the factors - such as HPRs - that support them. Particularly enigmatic are the heavily recombining mitogenomes of *T. gondii* and other members of the Sarcocystidae (Namasivayam *et al*., 2021; Tetzlaff *et al*., 2024). Transcript production in *T. gondii* mitochondria displays unique features, with rRNA fragments generated from recombination sites frequently overlapping in sense or antisense orientation with other rRNAs or with mRNAs (Tetzlaff *et al*., 2024; Shikha *et al*., 2025). The correct processing, stabilization, and ribosomal incorporation of such overlapping transcripts likely requires specialized mechanisms, potentially mediated by a subset of Sarcocystidae-specific HPR proteins.

### mRSiC as a coccidian-specific RNA stabilizer

We selected *T. gondii* mRSiC as a representative of the lineage-specific subset of HPR proteins for functional characterization. mRSiC is highly conserved among Toxoplasmatinae and has identifiable orthologs in *S. neurona* and *C. cayetanensis*, albeit with lower overall sequence conservation. As predicted for most HPR proteins (Hillebrand *et al*., 2018), mRSiC localizes to the *T. gondii* mitochondrion and contributes to parasite proliferation. Furthermore, our analyses demonstrated that mRSiC is required for the biogenesis and/or stability of ETC complexes containing mitochondrially encoded subunits, in particular Complex IV, suggesting that mRSiC facilitates the production of these subunits within mitochondria.

Monitoring the mitochondrial steady-state transcriptome during progressive mRSiC depletion revealed that mRSiC is specifically required for the accumulation or stability of two mitochondrial sRNAs, RNA33_1 and RNA42. Our data show that even moderately reduced mRSiC levels are limiting for preserving the steady-state levels of RNA33_1 and RNA42. We did not directly assess mitochondrial transcription and therefore cannot fully exclude that the decrease in RNA33_1 and RNA42 results from the involvement of mRSiC in transcription. However, given that primary transcripts in apicomplexan mitochondria are generated as long polycistronic precursors instead of monocistronic units (Ji *et al*., 1996; Namasivayam *et al*., 2021) and that RBPs predominantly function in post-transcriptional processes (Rackham, Mercer and Filipovska, 2012; Hammani *et al*., 2014; Rovira and Smith, 2019), mRSiC is more likely to act on the level of RNA stability rather than RNA production.

RNA33_1 represents a 24-nt subfragment that coincides with the 5’ end of the SSU rRNA fragment RNA33, which is itself severely reduced upon mRSiC loss. RNA33_1 could represent an individual mitochondrial sRNA of as-yet unknown function; however, its size, precise 5’ termini location and rapid decline upon mRSiC loss closely match the characteristics described for RBP-protected footprints (Ji *et al*., 2016; Ruwe *et al*., 2016). We therefore propose that RNA33_1 corresponds to a footprint generated by binding of an RBP, most parsimoniously mRSiC, to the RNA33 5’ end. RBP-derived footprints have been particularly well characterized for PPR proteins, helical repeat proteins abundant in plant organelles and structurally related to HPRs (Ruwe *et al*., 2016; Hillebrand *et al*., 2018). PPRs display high target specificity, typically binding individual RNAs or a small subset, with each repeat recognizing one nucleotide (Barkan *et al*., 2012). Strikingly, we observed that RNA33 and RNA42 exhibit perfectly overlapping sequence features at their 5’ ends that are not found in any of the other *T. gondii* mitochondrial transcripts. It is conceivable that the 7 HPR repeats of mRSiC may specifically bind these shared sequence elements at the 5’ ends of RNA33 and RNA42, further supporting the theory that RNA33_1 is a footprint that accumulates through the binding of mRSiC to the 5’ end of RNA33. Other helical hairpin protein regions of mRSiC not captured by our alveolate HPR profile may additionally contribute to RNA binding. However, despite these compelling *in silico* indications, future studies employing techniques such as enhanced crosslinking and immunoprecipitation (eCLIP) (Van Nostrand *et al*., 2016) are required to identify the mRSiC targets and precise binding sites experimentally.

How might destabilization of RNA33 and RNA42 in mRSiC-depleted mutants compromise ETC integrity and parasite fitness? Since RNA42 was newly identified in this study, its function remains to be elucidated. RNA42 is complementary to *coxI* mRNA and could potentially act by base-pairing (Pelechano and Steinmetz, 2013). However, there are only moderate changes in *coxI* levels during late mRSiC knockdown stages making this scenario less likely. Alternatively, RNA42 could correspond to one of the unassigned leftover rRNA fragments (ulr) whose sequence could not be resolved (Wang *et al*., 2024), possibly ulr19 of the SSU, based on its comparable length.

In the case of RNA33, a function as an rRNA fragment in the small subunit of the ribosome has been demonstrated (Wang *et al*., 2024). Thus, mRSiC could indirectly support mitoribosome biogenesis by protecting RNA33 from nucleolytic decay. Speculatively, mRSiC, might have an additional role in directly facilitating integration of RNA33 into the assembling ribosome. The assembly of the *T. gondii* mitoribosome from more than 40 rRNA fragments is proposed to be an elaborate and highly coordinated process (Wang *et al*., 2024; Shikha *et al*., 2025). Helical repeat proteins from plants, human and Trypanosomes have been suggested to be part of the mitoribosome assembly process (Kleinknecht *et al*., 2014; Antonicka and Shoubridge, 2015; Saurer *et al*., 2019; Méteignier *et al*., 2021). For instance, the PPR protein KRIPP2 binds to the 5’ end of 9S rRNA in Trypanosome mitochondria, facilitating its coordinated incorporation into the ribosome and KRIPP2 depletion is associated with destabilization of the 9S rRNA (Saurer *et al*., 2019). By analogy, transient interaction of mRSiC with the 5’ ends of RNA33 may promote assembly of this RNA into the *T. gondii* mitoribosome. In this scenario, the shorter RNA33_1 is just a degradation product of RNA33 protected by mRSiC. That the mitoribosome component RNA33 is more stable than RNA33_1 after mRSiC depletion might indicate that there are other routes into the mitoribosome independent of mRSiC. Detailed studies of mitoribosome assembly in mRSiC mutants are needed to test this hypothesis.

Irrespective of the underlying mechanisms, the strong decrease of RNA33 after loss of mRSiC likely impairs biogenesis and/or stability of the *T. gondii* mitoribosome, which in turn leads to compromised ribosome function. This consequently results in disturbed synthesis of the mitochondrially encoded ETC subunits and impaired parasite proliferation as observed in mRSiC-depleted mutants. The growth and ETC Complex IV defects observed upon mRSiC loss are comparatively mild relative to the severe phenotypes reported for mutants lacking mitoribosomal proteins (Lacombe *et al*., 2019; Shikha et al., 2022; Wang *et al*., 2024). This suggests that mitochondrial protein biosynthesis is impaired but not completely abolished after mRSiC depletion. Because RNA33 occupies a peripheral region of the SSU body (Wang *et al*., 2024), attenuated assembly within this region may not completely block ribosome biogenesis. Instead, partially functional ribosomes may still be produced, sufficient to maintain residual mitochondrial activity and parasite proliferation. In this context, reduced abundance of most of the remaining rRNA fragments at later stages of mRSiC depletion likely reflects pleiotropic effects arising from perturbed mitoribosome assembly or more generally from aberrant mitochondrial function.

Neither RNA33 nor RNA42 has been identified in the rRNA fragment repertoires reported for other alveolate lineages (Feagin *et al*., 2012; Tetzlaff *et al*., 2024), indicating that both RNAs are most likely specific to coccidians. This lineage-restricted occurrence is consistent with the confined distribution of mRSiC to coccidian parasites and suggests that mRSiC and its RNA targets may have evolved in parallel. Such a co-evolutionary scenario supports the idea that unusual mitochondrial gene expression systems in myzozoans can drive the emergence of novel, lineage-specific RNA-binding proteins.

## Materials and Methods

### Phylogenetic analysis

Predicted proteomes from 57 species were obtained from publicly available reference datasets, spanning apicomplexans, chromerids, dinoflagellates, perkinsids, ciliates, and a chlorophyte outgroup. To obtain an alveolate-specific consensus for heptatricopeptide repeat (HPR) proteins, we curated the experimentally supported HPR candidates reported by Hillebrand et al. (2018). These sequences, representing apicomplexans and additional alveolate lineages, were aligned using MAFFT v7 with the L-INS-i algorithm and otherwise default parameters (Katoh *et al*., 2005). Alignments were manually inspected to remove poorly aligned regions and low-quality sequences. A profile hidden Markov model (HMM) was built using HMMER v3.3.2 (Eddy, 2011) (hmmbuild), applying a minimum column occupancy threshold (--symfrac) of 0.7 to exclude sparsely populated alignment positions. The resulting model encodes a 37–amino-acid consensus repeat corresponding to a predicted helix–turn–helix (HTH) helical hairpin, characteristic of OPR/HPR-like repeat families. Each proteome was scanned using hmmsearch (HMMER v3.3.2), retaining both sequence-level and domain-level matches. Searches were performed using permissive significance thresholds (sequence E-value ≤ 10; domain E-value ≤ 10) to allow detection of divergent or partially degenerated repeat instances. All HPR-containing proteins were clustered into orthogroups using OrthoFinder v2.5 (Emms and Kelly, 2015). Clustering was based on sequence similarity within the HPR repeat regions, ensuring that orthogroups reflect evolutionary relationships of the repeat domain rather than unrelated non-repeat regions. OrthoFinder was run using DIAMOND for similarity searches and default settings for clustering. For each *T. gondii* HPR protein, its orthogroup distribution across the 57-species dataset was used to infer the most recent common ancestor (MRCA) of that group. MRCA assignments were mapped onto a schematic species tree reflecting consensus phylogenetic relationships (Stoeck *et al*., 2007; Hampl *et al*., 2009; Saffo *et al*., 2010; He *et al*., 2016; Janouškovec *et al*., 2017; Toscani Field *et al*., 2018; Kwong *et al*., 2021; Salomaki *et al*., 2021). This tree serves as a topological reference only and does not represent a new phylogenetic reconstruction. Orthologs of *T. gondii* TGME49_237530 (mRSiC) were retrieved from OrthoFinder orthogroups. Full-length sequences were aligned using MAFFT L-INS-i and automatically trimmed with trimAl. A maximum-likelihood phylogeny was inferred using IQ-TREE v3 (Wong *et al*., 2025), with model selection performed by ModelFinder and nodal support estimated using 1,000 ultrafast bootstrap replicates.

### Motif structural analysis and AlphaFold structural predictions

All extracted HPR motif windows were aligned and analyzed using a Python implementation of Chou–Fasman secondary-structure propensity scoring. For each alignment position, fractional α-helix, β-strand, and turn probabilities were calculated, revealing a characteristic pattern of two α-helical segments separated by a short turn, consistent with an HTH fold. Representative full-length HPR proteins, including *T. gondii* mRSiC, were structurally predicted using AlphaFold2

### Parasite culture

*T. gondii* parasites were cultured in human foreskin fibroblasts as described before (Jacot *et al*., 2020; Tetzlaff *et al*., 2024). When required, anhydrotetracycline (ATc) was added to a final concentration of 0.5 µg/ml. For harvesting, extracellular cultures were filtered through a 3 μm pore size polycarbonate filter, collected by centrifugation at 1500 × *g* for 10 min, and washed once in ice-cold Dulbecco’s phosphate-buffered saline (PBS) (Capricorn Scientific).

### Generation of genetically modified parasites

All transgenic parasite lines were generated using CRISPR/Cas9 as previously described (Shen *et al*., 2014). Parasites of the strain RH TATiΔ*ku80* (Sheiner *et al*., 2011) served as parental line. Single guide RNAs (sgRNAs) were selected using CHOPCHOP (Labun *et al*., 2019), https://chopchop.cbu.uib.no/). Primers used are listed in Table S1.

For C-terminal tagging of mRSiC (TGME49 237530), a sgRNA targeting the 3’ end of mRSiC was introduced into pSAG1::Cas9-U6::sgUPRT (Addgene plasmid # 54467; (Shen *et al*., 2014) by using the Q5-site directed mutagenesis kit (NEB), following the manufacturer’s instructions. A short glycine linker followed by a 3x FLAG tag sequence was integrated into pCas9-CAT (Addgene plasmid # 54467; (Sidik *et al*., 2016) and the resulting linker/3xFLAG/CAT cassette amplified using Q5 polymerase (NEB) and primers containing 50 bp overhangs homologous to the either upstream or downstream regions of the mRSiC stop codon. The amplified donor construct was co-transfected together with the pSAG1::Cas9-U6::sgUPRT vector encoding the sgRNA and Cas9-GFP into RH TATiΔ*ku80* parasites. Transgenic parasites were selected with chloramphenicol and cloned by limiting dilution as previously described (Roos *et al*., 1994). Genotyping of single clones was performed as described in (Piro, Carruthers and Di Cristina, 2020) and correct integration of the 3xFLAG tag verified by Sanger Sequencing.

A mRSiC conditional knockdown strain (termed rmRSiC-FLAG) was generated using the anhydrotetracycline (ATc)-regulatable TET-Off system, as described previously (Sheiner *et al*., 2011; Katris *et al*., 2014). The pSAG1::Cas9-U6::sgUPRT transfection vector was prepared as described above, with a sgRNA targeting the 5’ region of mRSiC near the start codon. The donor construct was amplified from the vector pPR2-HA3-Floxed-DHFR, a derivative of pPR2-HA3 (Katris *et al*., 2014), using primers with homology arms corresponding to the regions up-and downstream of the mRSiC start codon. Transfection, limiting dilution, and genotyping were performed as described above, with the mRSiC-FLAG strain serving as the parental line and selection performed with pyrimethamine.

Parasite lines expressing tobacco etch virus (TEV) protease cleavage site–HA-tagged versions of ETC subunits were derived from rmRSiC-FLAG. Transfection constructs were generated following the same principles used for C-terminal tagging of mRSiC. Existing sgRNA-expression plasmids were used for TgSdhB, TgMPPa and TgCox2a (Seidi *et al*., 2018; Hayward *et al*., 2021; Leonard *et al*., 2023) and parasite lines were generated as described in (Hayward *et al*., 2021). Three days post-transfection, GFP-positive parasites were isolated and cloned by fluorescence-activated cell sorting (FACS) and subsequently genotyped as described above.

### Plaque assays

Plaque assays were conducted as previously described (Jacot *et al*., 2020). Confluent T25 host cell flasks were infected with 400 parasites, incubated for 7 days and afterwards stained with crystal violet. Plaque area quantification was automated using the following custom pipeline. Images were converted to grayscale and denoised using Gaussian and optional median blurs, followed by Contrast Limited Adaptive Histogram Equalization (CLAHE) to enhance contrast. Following global thresholding and morphological closing to refine plaque boundaries, contours were detected with OpenCV, and plaques within a defined size range were retained for area measurement.

### Immunofluorescence/Fluorescence microscopy assays

Immunofluorescence assays (IFAs) were performed as described by (van Dooren *et al*., 2008). We used the following primary antibodies: rat anti-HA High Affinity clone 3F10 (1:500, Roche, ROAHAHA), mouse anti-FLAG M2 (1:250, Sigma, F3165), and rabbit anti-Tom40 (1:2,000, (van Dooren *et al*., 2016). Secondary antibodies used were goat anti-rat Alexa Fluor 488 (1:500, Thermo Fisher Scientific, A-11006), goat anti-mouse Alexa Fluor 488 (1:500, Thermo Fisher Scientific, A-11029), and goat anti-rabbit Alexa Fluor 594 (1:500, Thermo Fisher Scientific, A11012). Images were acquired on a DeltaVision Elite deconvolution microscope (GE Healthcare) with a 100X UPlanSApo objective lens (NA 1.40). They were deconvolved using SoftWoRx Suite 2.0 software and contrast and brightness were adjusted in FIJI/ImageJ. Images were then processed with Adobe Illustrator.

### SDS-PAGE, BN-PAGE and immunoblotting

SDS-PAGE and immunoblotting were performed as described by (Tetzlaff *et al*., 2024). Per lane, 2.5×10^6^ parasites were loaded. Blue native (BN)-PAGE followed the protocol by (van Dooren *et al*., 2016). Parasites were solubilized in either 0.5% digitonin (for Complexes II and V) or 1% Triton X-100 (for Complexes III and IV). Blots were probed with the following primary antibodies: mouse anti-FLAG M2 (1:500–1:2000, Sigma, F3165), rat anti-HA High Affinity clone 3F10 (1:500–1:1000, Roche, ROAHAHA), and rabbit anti-Tom40 (1:2000, (van Dooren *et al*., 2016). Secondary antibodies conjugated to horseradish peroxidase (HRP) were goat anti-mouse IgG H&L (1:10,000, abcam, ab205719), rabbit anti-rat IgG H&L (1:5000, abcam, ab6734), and goat anti-rabbit IgG H&L (1:10,000, abcam, ab205718).

### Small RNA Sequencing

Freshly harvested parasites corresponding to one extracellular T175 flask culture were subjected to 0.05% digitonin treatment for crude organellar enrichment as described previously (Tetzlaff *et al*., 2024). The pellet obtained after treatment and centrifugation was resuspended in TRIzol Reagent (Invitrogen) and stored at −80°C. RNA was isolated using the Monarch Total RNA Miniprep Kit (New England BioLabs) according to the manufacturer’s protocol for TRIzol extracted samples. Afterwards, RNA samples were subjected to a customized rRNA depletion procedure following the protocol published by (Thompson, *et al*., 2020). In total, 36 biotinylated probes were used to target the *T. gondii* cytosolic 28S, 18S, 5.8S, and 5S rRNA, as well as the apicoplast LSU and SSU rRNAs (oligonucleotide sequences are listed in Table S1). Probe selection was additionally revised to exclude cross-hybridization with mitochondrial rRNA fragments. Next, 175 ng of rRNA-depleted RNA was used for the preparation of small RNA sequencing libraries using the NEBNext Multiplex Small RNA Library Prep Set for Illumina Kit (New England BioLabs), following the manufacturer’s instructions. Libraries were sequenced on an Illumina NovaSeq platform (150 bp read length, paired end) at Azenta Life Sciences.

### Identification of novel sRNA features

Read start and end positions were aggregated at single-nucleotide resolution. For each sample, signal-to-noise ratios (SNRs) for read starts and ends were calculated at position x as the ratio between the observed count and the median count within a ±25-nt window. To reduce technical variability, only the strongest peak within a ±3-nt window was retained. Candidate feature boundaries were required to show ≥100 reads and an SNR ≥50 in all three biological replicates of at least one condition. Final feature boundaries were defined by manual inspection.

### Small RNA Sequencing data analysis

Illumina reads were quality filtered and adapter- and poly(A)-trimmed using cutadapt (v4.6; -q 20 -m 15 --trim-n --poly-a). Trimmed reads were mapped to a custom mitochondrial pseudo-genome and nuclear/apicoplast decoy references (bowtie v1.3.1; ungapped alignment, ≤1 mismatch, multimapping enabled). Read origins for reads mapping to both references were determined by a score-based comparison: assignments were first based on mismatch count, with ties resolved by mapping multiplicity, favouring the reference with fewer distinct mapping loci. To minimise ambiguous counting, reads were assigned only to features they covered by at least 80% and counted exclusively toward the feature with the largest overlap using featureCounts (v2.0.6). Longer features for which the 80% criterion could not be satisfied were quantified using the largest-overlap rule alone. Only sense-strand alignments were used for quantification, and read identifiers were retained for downstream poly(A)-tail analyses. Decoy-exclusive reads were quantified for all non-rRNA features. For normalisation, decoy-mapped features were screened for correlation with the ATc-induced perturbation, and features showing low correlation across the treatment series were selected as experiment-specific reference features; nuclear rRNAs were excluded. Mitochondrial fragment counts were analysed using edgeR (v4.4.2), with negative-binomial dispersion estimated in the standard framework, but library sizes overridden by the total counts of the selected reference feature set. Differential expression was assessed using quasi-likelihood F-tests (glmQLFTest). For sense-strand mitochondrial reads, poly(A)-tail length was estimated from terminal homopolymeric A/T stretches remaining after adapter trimming. Read-level tail lengths were aggregated per fragment and condition. Differences in poly(A)-tail length distributions were quantified using the Earth Mover’s Distance (EMD). Statistical significance was assessed by permutation testing (1,000 permutations), in which tail lengths were randomly reassigned between conditions to generate a null distribution. To correct for background noise, observed EMD values were additionally required to exceed fragment- and condition-specific background variability estimated from technical replicate comparisons. All code, reference files, and the final pipeline are available at https://github.com/Kovox91/toxo_srna_pipeline; intermediate files supporting annotation and fragment-boundary curation are available upon request.

### Small RNA gel blot hybridization

Small RNA gel blot analysis of mitochondrial rRNA fragments was performed as described in (Tetzlaff *et al*., 2024). Sequences of antisense oligonucleotide probes used are listed in Table S1.

### mRNA gel blot hybridization

RNA gel blot hybridization for the detection of mitochondrial mRNAs was performed as previously described (Kupsch *et al*., 2012) with minor modifications. 5 µg of total RNA were loaded per lane and separated by electrophoresis. Primers used to amplify PCR products serving as templates for RNA probe generation are listed in Table S1. Prehybridization and Hybridization were performed at 64°C. Afterwards, membranes were washed once with 2× SSC, 0.1% SDS and once with 1× SSC, 0.1% SDS at 64°C. Between consecutive hybridizations, probes were removed by three rounds of adding 0.5% boiling SDS followed by incubation for 30 minutes 70 °C.

### Small RNA sequence motif search

Shared sequence motifs between RNA33 and RNA42 were identified using the MEME suite software (Bailey *et al*., 2015) and Clustal Omega (Sievers *et al*., 2011). The occurrence of the output motif within the *T. gondii* mitochondrial genome was assessed using FIMO (Grant, Bailey and Noble, 2011).

## Data availability

RNA sequencing data generated in this study have been deposited in the NCBI BioProject database (https://www.ncbi.nlm.nih.gov/bioproject/) under accession number PRJNA1380901.

## Acknowledgments

We sincerely thank Luisa Bernhardt for establishing the rRNA depletion procedure and Edwin Tjhin, formerly of the van Dooren Group, for preparing the sgRNA-expression plasmids for Atpβ tagging.

## Author’s contribution

S.T., C.S.L, G.v D., and R.W. designed and supervised the research; N.D., Z.G., B.A.B., A.H., A.P., and S.T. performed the experiments and interpreted the data; N.D., S.M., and V.F. analyzed datasets and interpreted the data; S.T. wrote the manuscript with contributions from all authors.

