## Supplemental Figures for "Helical Repeat Protein mRSiC is Required for rRNA Fragment Accumulation in *T. gondii* Mitochondria"

### Drakoulis et al. – Supplemental Figures

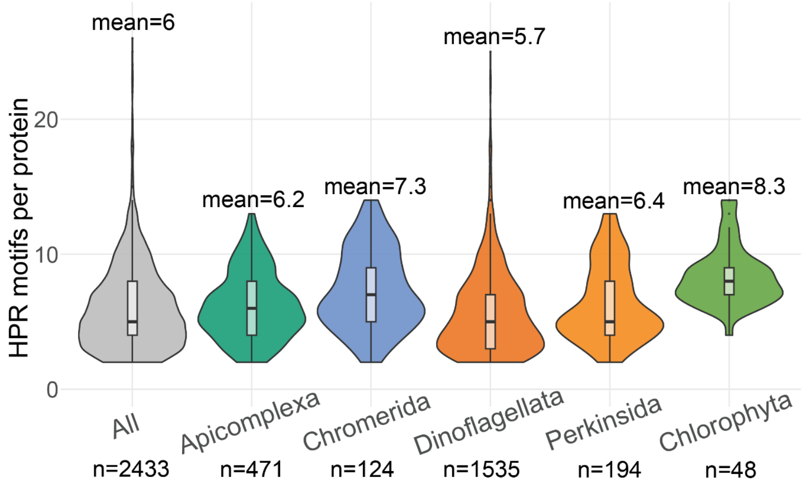


**Figure S1. HPR repeat counts in different lineages.**
HPR motif repeat counts across major lineages included in the HMM search. For each HPR-positive protein (≥2 tandem repeats), the total number of repeats was quantified and visualized per clade using violin/box plots.

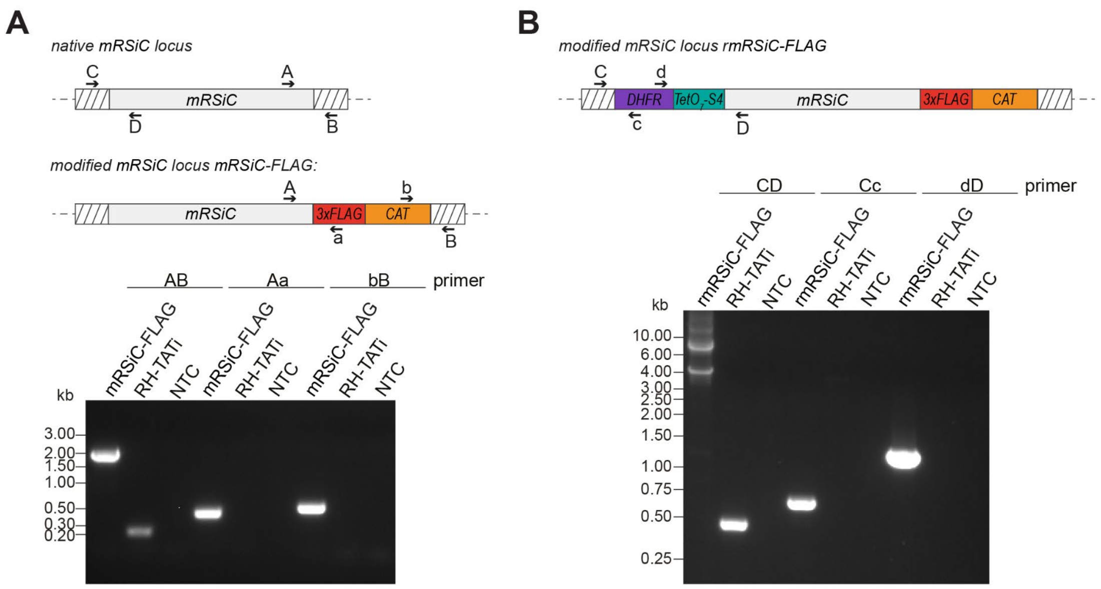


**Figure S2. Tagging and knock-down approach for mRSiC (TGME49 237530).**

**(A)** Top panel: Schematic representation of the native *mRSiC* locus (top) and the locus after insertion of a triple FLAG (*3xFLAG*) epitope tag at the 3′ end of the open reading frame together with the chloramphenicol acetyltransferase (*CAT*) selectable marker (bottom). Primer binding sites used to test integration are indicated by black arrows. Bottom panel: Genotyping PCR analysis of genomic DNA from mRSiC-FLAG parasites using the primer pairs shown in the schematics above. RH-TATi Δku80 genomic DNA and no-template controls (NTC) were included as controls. The results confirm integration of the *3xFLAG* epitope tag at the 3’ end of mRSiC.

**(B)** Top panel: Schematic representation of the *mRSiC* locus after insertion of an anhydrotetracycline (ATc)-regulatable TetO7-Sag4 (TetO7-S4) promoter, along with a dihydrofolate reductase (*DHFR*) selectable marker cassette, upstream of the start codon of the *mRSiC* gene, which had previously been modified to express a C-terminal 3xFLAG tag (see A). Primer binding sites used to screen for promoter integration are shown by black arrows. Bottom panel: Genotyping PCR analysis of genomic DNA from rmRSiC-FLAG parasites using the primers shown above to assess the integration of the promoter cassette. RH-TATi Δku80 genomic DNA and no-template controls (NTC) were included as controls. Results indicate successful integration of the TetO7-Sag4 promoter upstream of the mRSiC open reading frame.

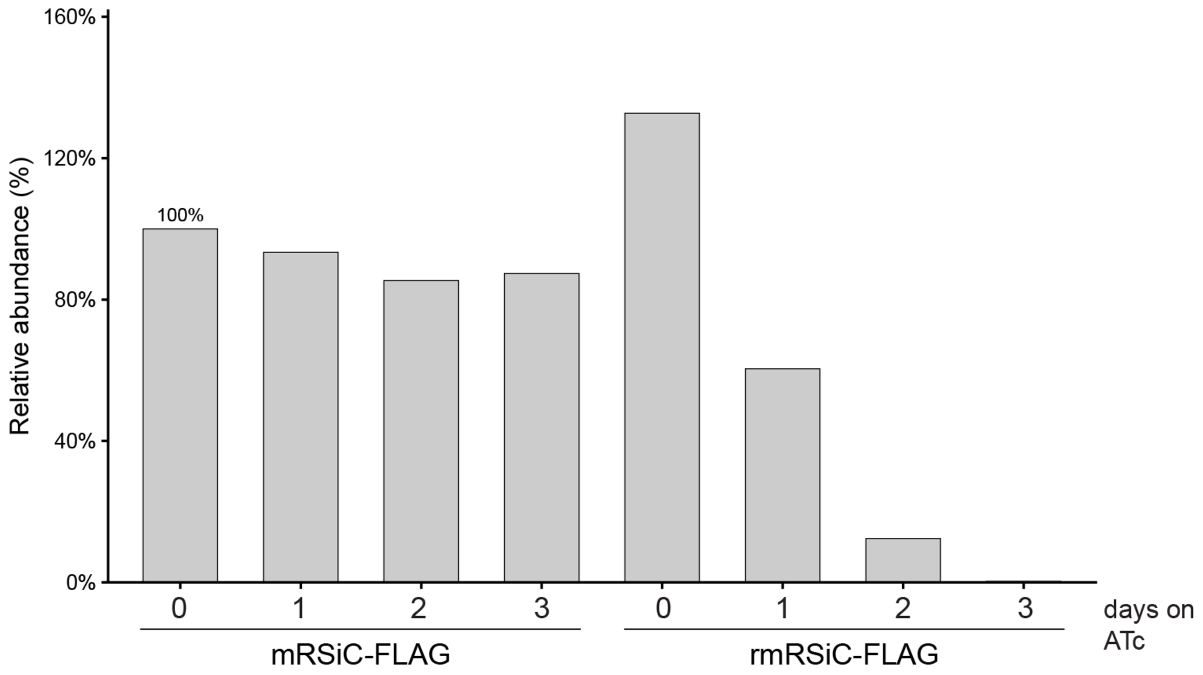


**Figure S3. Quantification of immunoblot signals assessing mRSiC knockdown efficiency.** mRSiC-FLAG signals from the immunoblot shown in 3A were normalized to Tom40 and visualized relative to the signal intensity in untreated parental mRSiC-FLAG parasites.

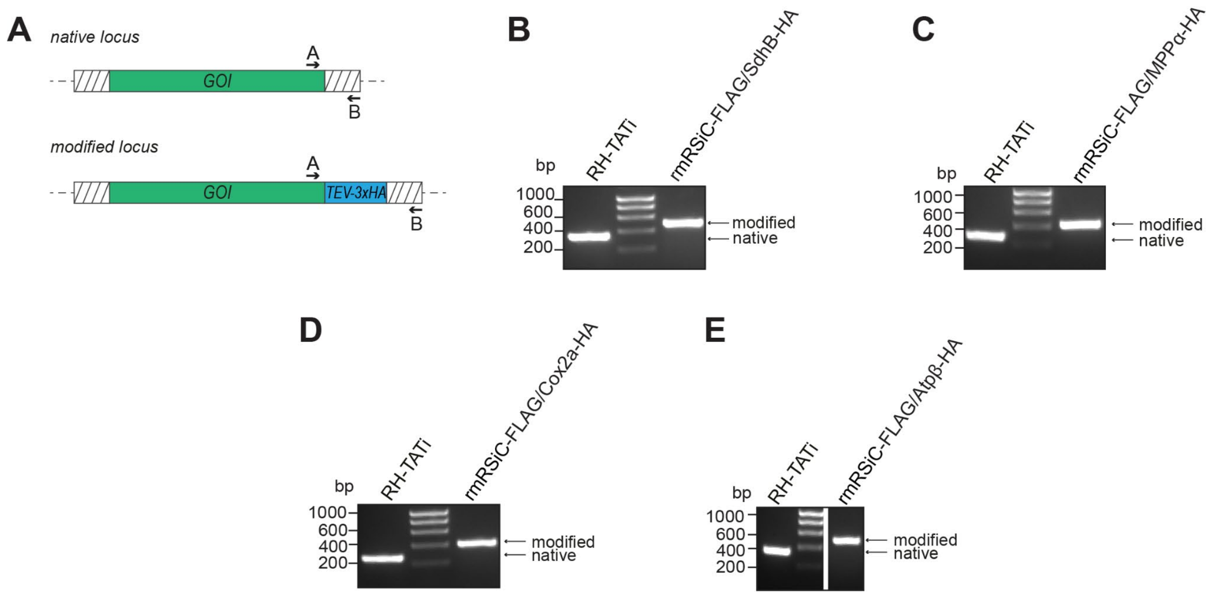


**Figure S4. HA-tagging of mitochondrial ETC subunits in the rmRSiC-FLAG strain.**

**(A)** Schematic illustrating the integration of a TEV-3xHA epitope tag at the 3’ end of genes encoding ETC subunits in the rmRSiC-FLAG background. Primer binding sites used to assess correct integration are indicated by black arrows. Selection of transfected parasites was performed by fluorescence-activated cell sorting (FACS) based on GFP fluorescence from the co-transfected Cas9-GFP expression vector. *GOI*, gene of interest

**(B)** Genotyping PCR analysis of genomic DNA from rmRSiC-FLAG/SdhB-HA using the primers A and B indicated in the schematic above. Genomic DNA from RH-TATi Δku80 parasites was included as a control. The results verify integration of the 3xHA tag at the 3’ end of SdhB in the rmRSiC-FLAG/SdhB-HA strain.

**(C-E)** As in panel B, PCR analyses confirming the generation of the strains rmRSiC-FLAG/MPPa-HA (C), rmRSiC-FLAG/Cox2a-HA (D), and rmRSiC-FLAG/Atpβ-HA (E). White lines indicate where lanes irrelevant to this analysis were removed.

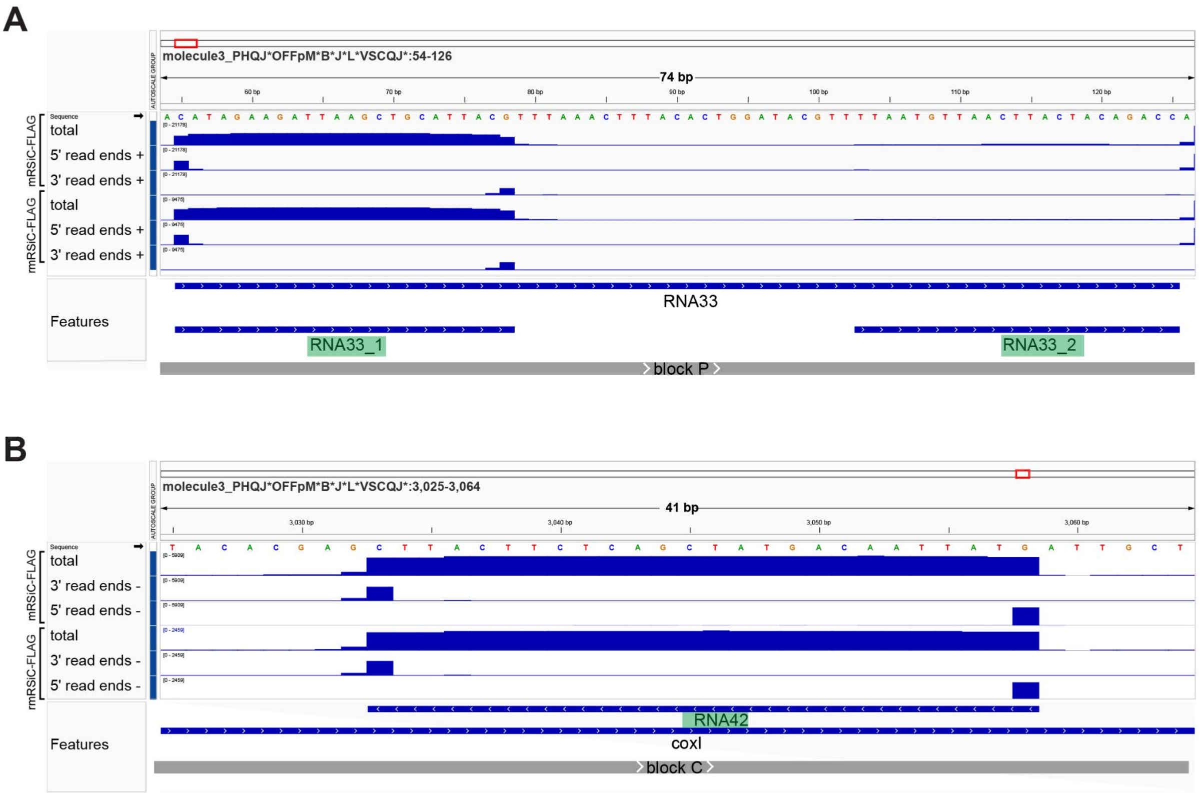


**Figure S5. Annotation of additional mitochondrial features based on sRNA-sequencing of rRNA-depleted samples.**

**(A)** Excerpts of coverage plots showing read depth at the RNA33 locus in sRNA-sequencing samples from untreated mRSiC-FLAG and rmRSiC-FLAG parasites. Coverage is shown for full-length reads (total) as well as for 5'- and 3' read ends. "−" and "+" indicate whether coverage is shown for the forward or reverse strand, respectively. Mitochondrial genomic sequence blocks are shown in gray. Annotated features are displayed in blue below, with features not annotated previously (Wang et al., 2024) highlighted in green.

**(B)** Same as in (A), but for the RNA42 locus.

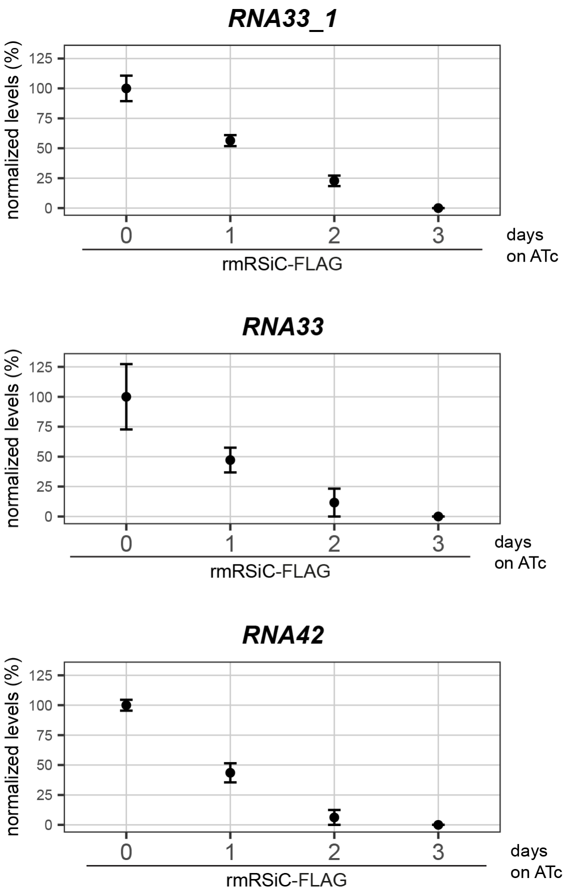


**Figure S6. Small RNA gel blot quantification examining mitochondrial sRNA levels in mRSiC knockdown parasites.**

Quantification of signal intensities from sRNA gel blot hybridizations shown in Fig 6D. Signals were normalized to the 5.8S rRNA signal and are visualized as linearly scaled absolute mean values ± SEM, with untreated samples (0 day ATc treatment) set to 100%, n=2

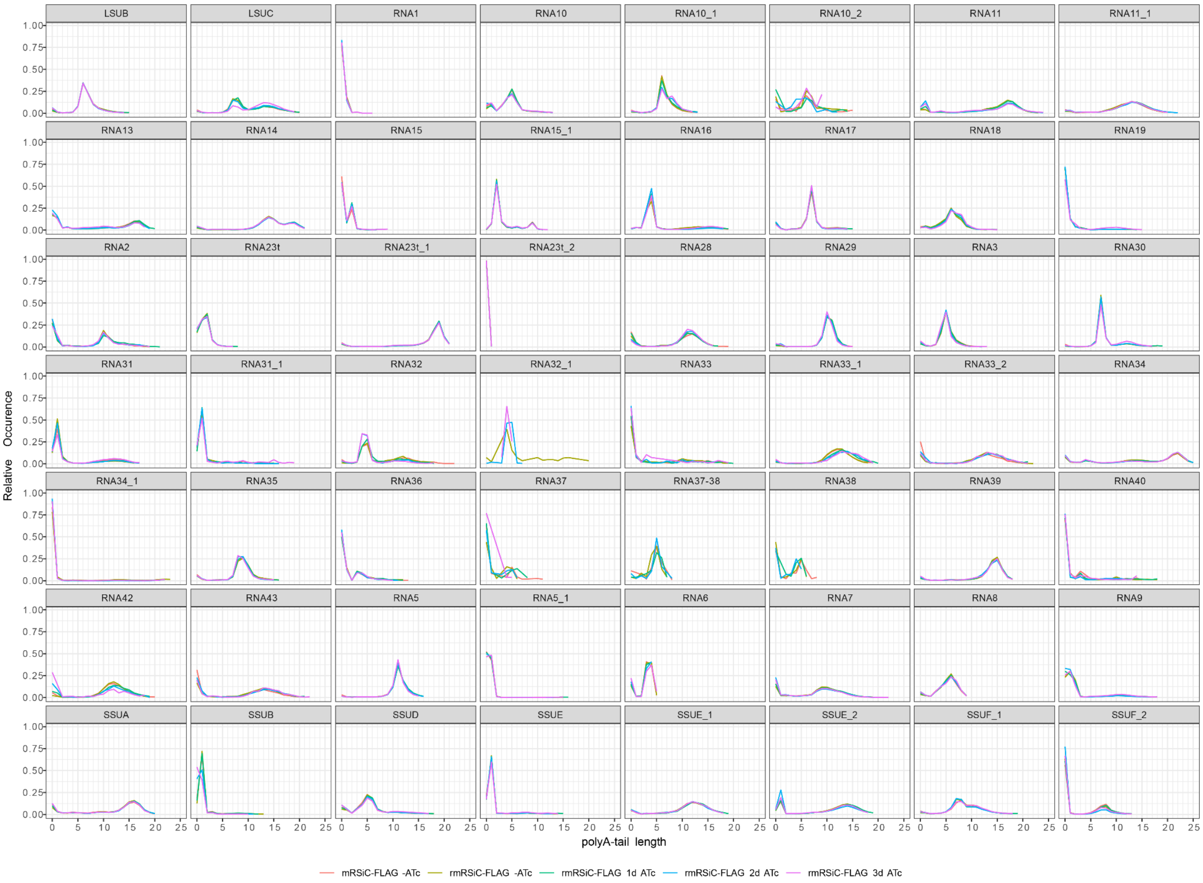


**Figure S7: Mean poly(A)-tail length distributions of mitochondrial transcripts across conditions, shown per gene (facets).**

Line plots represent average fragment-level poly(A)-tail profiles for each condition. Distributional differences were quantified using the Wasserstein (earth mover's) distance and assessed by permutation testing (1,000 permutations). Gene–condition combinations exceeding background variability (Wasserstein distance > background +1) and passing FDR correction (FDR < 0.05) are reported in Supplementary Table S4.

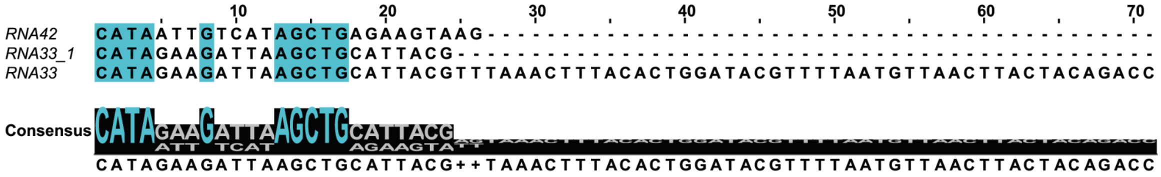


**Figure S8. RNA42 and RNA33/RNA33_1 and share sequence elements at their 5’ ends.**
Sequences of RNA42, RNA33_1 and RNA33 were aligned using Clustal Omega. Identical nucleotide positions between the sequences are shaded in turquoise. The automated consensus sequence is displayed below.
